# Genomic evidence for West Antarctic Ice Sheet collapse during the Last Interglacial

**DOI:** 10.1101/2023.01.29.525778

**Authors:** Sally C. Y. Lau, Nerida G. Wilson, Nicholas R. Golledge, Tim R. Naish, Phillip C. Watts, Catarina N. S. Silva, Ira R. Cooke, A. Louise Allcock, Felix C. Mark, Katrin Linse, Jan M. Strugnell

## Abstract

The marine-based West Antarctic Ice Sheet (WAIS) is considered vulnerable to irreversible collapse under future climate trajectories and its tipping point may even lie within the mitigated warming scenarios of 1.5–2 °C of the United Nations Paris Agreement. Knowledge of ice loss during similarly warm past climates, including the Last Interglacial, when global sea levels were 5–10 m higher than today, and global average temperatures of 0.5–1.5 °C warmer than preindustrial levels, could resolve this uncertainty. Here we show, using a panel of genome-wide, single nucleotide polymorphisms of a circum-Antarctic octopus, persistent, historic signals of gene flow only possible with complete WAIS collapse. Our results provide the first empirical evidence that the tipping point of WAIS loss could be reached even under stringent climate mitigation scenarios.

**One-Sentence Summary:** Historical gene flow in marine animals indicate the West Antarctic Ice Sheet collapsed during the Last Interglacial.

---

Climate change continues to cause unprecedented warming to the Earth system (1). The consequences of warming are leading to rapid changes in Antarctica, including Antarctic Ice Sheet mass loss, with global impacts (1). A major uncertainty in global mean sea level (GMSL) rise projections lies in the potential instability of the West Antarctic Ice Sheet (WAIS) (2). The marine-based WAIS has lost 159 ± 8 gigatons of ice mass per year between 1979–2017 (3), and will continue to be a major contributor to GMSL rise under all CO_2_ emission scenarios (4). It is unclear whether the WAIS is vulnerable to rapid ice loss or even full collapse, because of a poor understanding of future changes and processes that influence ice sheet dynamics (2). WAIS collapse could raise global sea level by ∼3.3–5 m (5, 6), with direct consequences that include human displacement and global loss of ecosystems in coastal areas (1).

It is well understood from geological reconstructions that there were interglacial peaks, referred to as super-interglacials, in periods of the Pleistocene that experienced warmer temperatures (+∼0.5–1.5 °C) and higher GMSL (up to +10 m) than present (4). These super-interglacials include Marine Isotope Stages (MIS) 31, 11 and 5e, which occurred at approximately ∼1.08–1.05 Ma, ∼424–395 ka and ∼129–116 ka, respectively (4). During MIS 31, the Southern Ocean sea surface temperature may have reached +5 ± 1.2 °C above present during summer months (7).

During MIS 11, global mean surface temperature (GMST) was 0.5 ± 1.6°C with GMSL 6–13 m higher than present, and similarly, during MIS 5e (the Last Interglacial), GMST was +0.5–1.5°C with GMSL 5–10 m higher than present (4). To date, there is no empirical evidence to indicate if the WAIS has completely collapsed at any time in the three million years since the Pliocene (8, 9). Inferring WAIS configurations during late Pleistocene super-interglacial periods could therefore inform the sensitivity of Antarctic ice-sheet response to climate change. So far, analyses of ice proximal marine drill core records show evidence of WAIS retreat during the late Pleistocene interglacials, but the exact timing (10) and extent (9, 11) of any WAIS collapse remain ambiguous. Existing ice sheet models have yielded conflicting WAIS reconstructions during these periods, ranging from no collapse (12), to partial (13) or full collapse (14, 15). Knowledge about how the WAIS was configured during super-interglacials in the geological past is urgently needed to constrain future sea-level rise projections (2). Novel approaches, such as population genomics, can serve as empirical proxies of past changes to the Antarctic Ice Sheet, detected via signals of historic gene flow among currently separated populations of marine organisms (16).

A complete past collapse of the WAIS would have opened the trans-West Antarctic seaways linking the present-day Weddell Sea (WS), Amundsen Sea (AS) and Ross Sea (RS) (16). Such seaways would have allowed marine benthic organisms to occupy and disperse across the opened straits, thus leaving genetic signatures of this past connectivity in the genomes of their descendent, extant populations (16) (hereafter seaway populations). As the WAIS reformed, these organisms would be isolated again within the WS, AS and RS basins, with any subsequent connectivity only possible around the continental margin. Although there is some support for existence of trans-Antarctic seaways based on species assemblage data at macro-evolutionary scales (17–20) or low-resolution genetic data (21–24), all these studies lack power and/or spatial coverage to distinguish between past dispersal via trans-West Antarctic seaways or from contemporary circumpolar ocean currents. Importantly, these previous studies cannot be used for accurate demographic modelling to identify the likely timing of any collapse of the WAIS.

Collection of benthic species from the Southern Ocean is logistically challenging and regions such as AS and East Antarctica (EA) are difficult to access. Existing samples are typically characterised by DNA degradation due to long term storage in collections at room temperature. Here, we used a target capture approach that sequenced genome-wide, single nucleotide polymorphism (SNP) data in the circum-Antarctic benthic octopus, *Pareledone turqueti*, incorporating rare samples from AS and EA, collected over 33 years. Our approach enabled a comprehensive sampling strategy to robustly date and test for the presence of past trans-West Antarctic seaways using biological data as proxies.

We sequenced genome-wide SNPs derived from double-digest restriction site-associated DNA (ddRAD) (25) loci from 96 *P. turqueti* individuals collected from around the Southern Ocean (Fig. 1A). The dataset presents a circum-Antarctic overview of the species genetic patterns, which record the contemporary connectivity driven by oceanic currents, mainly the Antarctic circumpolar current (ACC; clockwise) and the Antarctic Slope Current (ASC; counter-clockwise) (Fig. 1A, B), as well as any historical connectivity that would be associated with past trans-West Antarctic seaways. We used a reduced single-nucleotide polymorphisms (SNPs) dataset (one SNP per locus) to analyse population structure, which included 5,188 unlinked SNPs. Complementary analyses (*Structure*, *TreeMix*) suggest the population genomic variation of *P. turqueti* is characterised by geographically-structured populations across the Southern Ocean (Fig. 1C, fig. S1-S2). This makes it an appropriate species to test for the presence of trans-West Antarctic seaways as signals of historical WAIS collapse in highly admixed species would likely be masked by contemporary gene flow signatures (16). *Pareledone turqueti* can disperse via benthic crawling as adults and as hatchlings (inferred from large egg size [maximum oocyte length=19.8mm] and small number of eggs per clutch [n=22-66]) (26, 27). It has also been suggested that long-distance dispersal in *P. turqueti* may be achieved, at least occasionally, via adults or egg masses rafting on floating substrates, or that their benthic egg masses could become dislodged and disperse through the currents (23), although no direct evidence supports this yet.

**Fig. 1.**
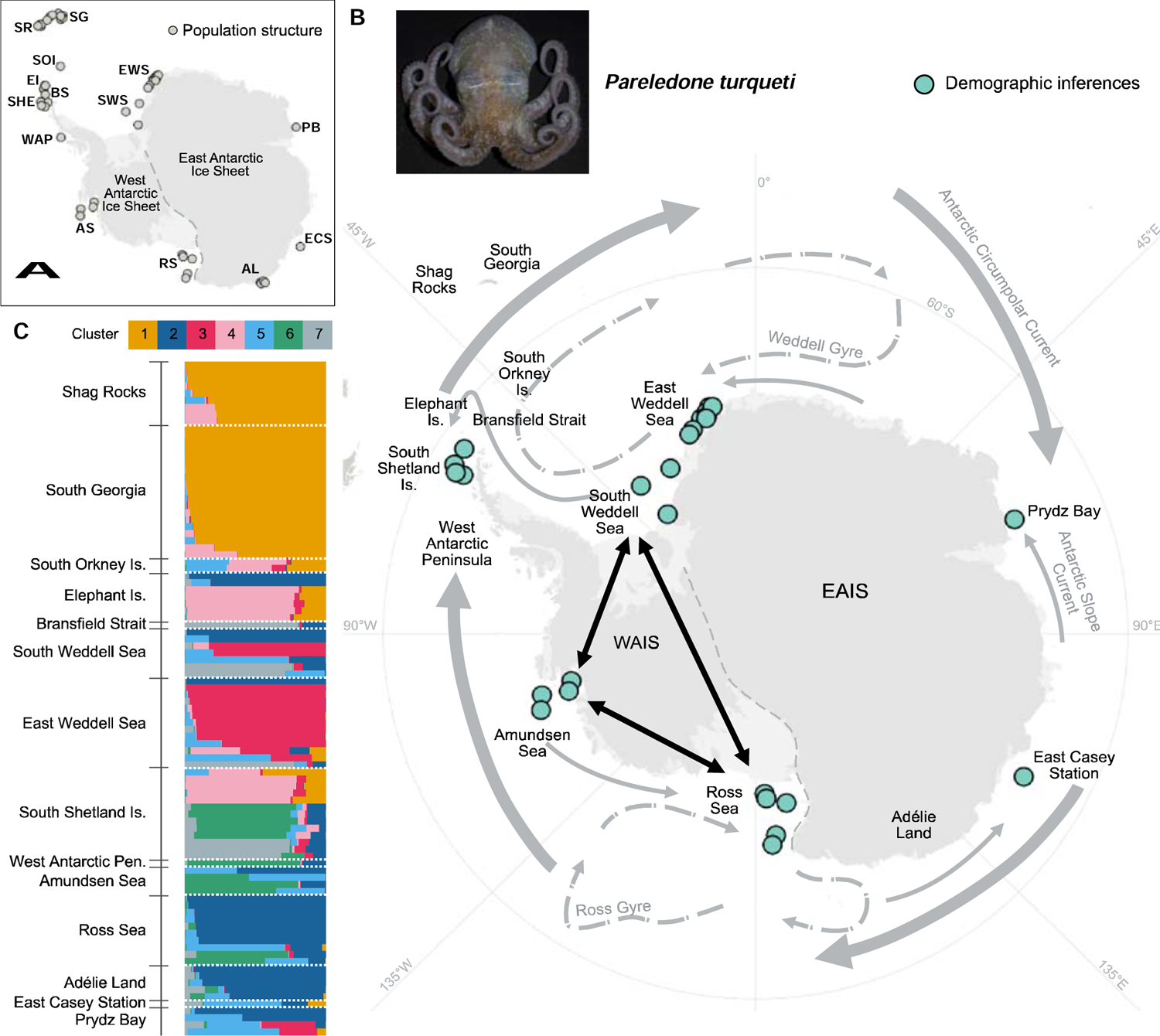
Sample locations of *Pareledone turqueti* with *Structure* analyses. (**A**) Samples used for analyses of population structure. Abbreviations: Shag Rocks, SR; South Georgia, SG; South Orkney Is., SOI; Elephant Is., EI; Bransfield Strait, BS; South Shetland Is., SHE; West Antarctic Peninsula, WAP; South- and East-Weddell Sea, S-, E-WS; Amundsen Sea, AS; Ross Sea, RS; Adélie Land, AL; East Casey Station, ECS; Prydz Bay, PB; West Antarctic Ice Sheet, WAIS; East Antarctic Ice Sheet, EAIS. (**B**) Samples used for admixture analyses and demographic modelling (collectively demographic inferences) to test for the existence of trans-West Antarctic seaways. Map includes the directionalities of the major contemporary circumpolar currents and regional currents in the Southern Ocean. Black arrows indicate connectivity pathways through trans-West Antarctic seaways that would result from WAIS collapse. Direct connectivity between WS-AS or AS-RS would indicate partial WAIS collapse, and direct connectivity between WS-AS-RS or WS-RS would indicate complete WAIS collapse. Photo credit *of P. turqueti* specimen: Elaina M. Jorgensen. (**C**) Clustering analysis using *Structure* inferred *K* = 7 for *P. turqueti* (5,188 SNPs dataset). Each horizontal bar represents an individual sample, bars are grouped by geographical locations, colours within each bar correspond to the proportion of each genetic cluster in the individual.

In *P. turqueti*, long-distance connectivity linking East and West Antarctica is detected across the Antarctic continental shelf and Antarctic islands (Fig. 1C), likely indicating dispersal that reflects contemporary circumpolar currents, as found in other Southern Ocean benthic taxa (28). If the genomic data of *P. turqueti* can only be explained by serial circumpolar colonisation around the Antarctic continent, then observed and expected heterozygosity would decrease as geographical distance increases, yet the observed data does not support this scenario (fig. S3, table S1). Admixture is also observed between individuals from RS and AS with some individuals from WS (Fig. 1C), indicating a potential signature of trans-West Antarctic seaways. Supporting this concept, population differentiation analysis shows limited genetic divergence between WS-RS relative to other localities that are adjacent to each other (fig. S4).

Focusing on populations most informative of whether the WAIS collapsed in the past, we first examined whether there was distinct admixture between WS-AS-RS with respect to South Shetland Islands (SHE) and East Antarctica (EA; including Prydz Bay and East Casey Station) samples using 120,857 SNPs (Fig. 1B). SHE and EA are known to be influenced by both the ACC and ASC (29), but are peripheral to the putative historical trans-West Antarctic connectivity; thus these are ideal locations that can separate patterns of present-day connectivity around the WAIS and East Antarctic Ice Sheet (EAIS) from persistent, historical signals of gene flow.

We examined allele frequency correlations across WS, AS and RS with respect to SHE and EA. The *D*-statistic (30) explores the patterns of allele sharing across four populations to test for evidence of admixture between populations of interest. The outgroup-ƒ_3_-statistic (31) explores the amount of derived allele frequency that is shared between pairs of populations relative to an outgroup population. The presence of admixture linked to trans-West Antarctic connectivity would result in high ƒ_3_ values, and evidence of excess allele sharing (*D*>0), between WS-AS-RS. In *P. turqueti*, the highest ƒ_3_ values are detected between AS–RS, followed by RS–EA and RS– WS (Fig. 2A); indicating recent common ancestry across seaway populations, as well as between RS and EA populations that are adjacent to each other. When SHE is treated as the sister lineage to AS/RS and WS (*D*(AS/RS, SHE, WS, outgroup)), there is excess allele sharing between AS/RS and WS (Fig. 2B). When EA is treated as sister lineage to AS/RS and WS (*D*(AS/RA, EA, WS, outgroup)), excess allele sharing is also observed between AS/RS and WS (Fig. 2B). If circumpolar currents (ASC and ACC) were the only factors that have influenced gene flow patterns in *P. turqueti*, then low ƒ_3_ values would be observed between WS-RS as they are situated on the opposite side of West Antarctica, and excess allele sharing (*D*) would also be observed between WS-SHE and WS-EA as they are geographically adjacent. However, these results confirmed that in *P. turqueti* there are unexpected and significant allele frequency correlations among AS-RS-WS, despite also considering the locations situated between them around the WAIS (SHE) and EAIS (EA). Such observed admixture patterns are congruent with historical seaway connectivity in a species that is characterised by geographically-structured populations.

**Fig. 2.**
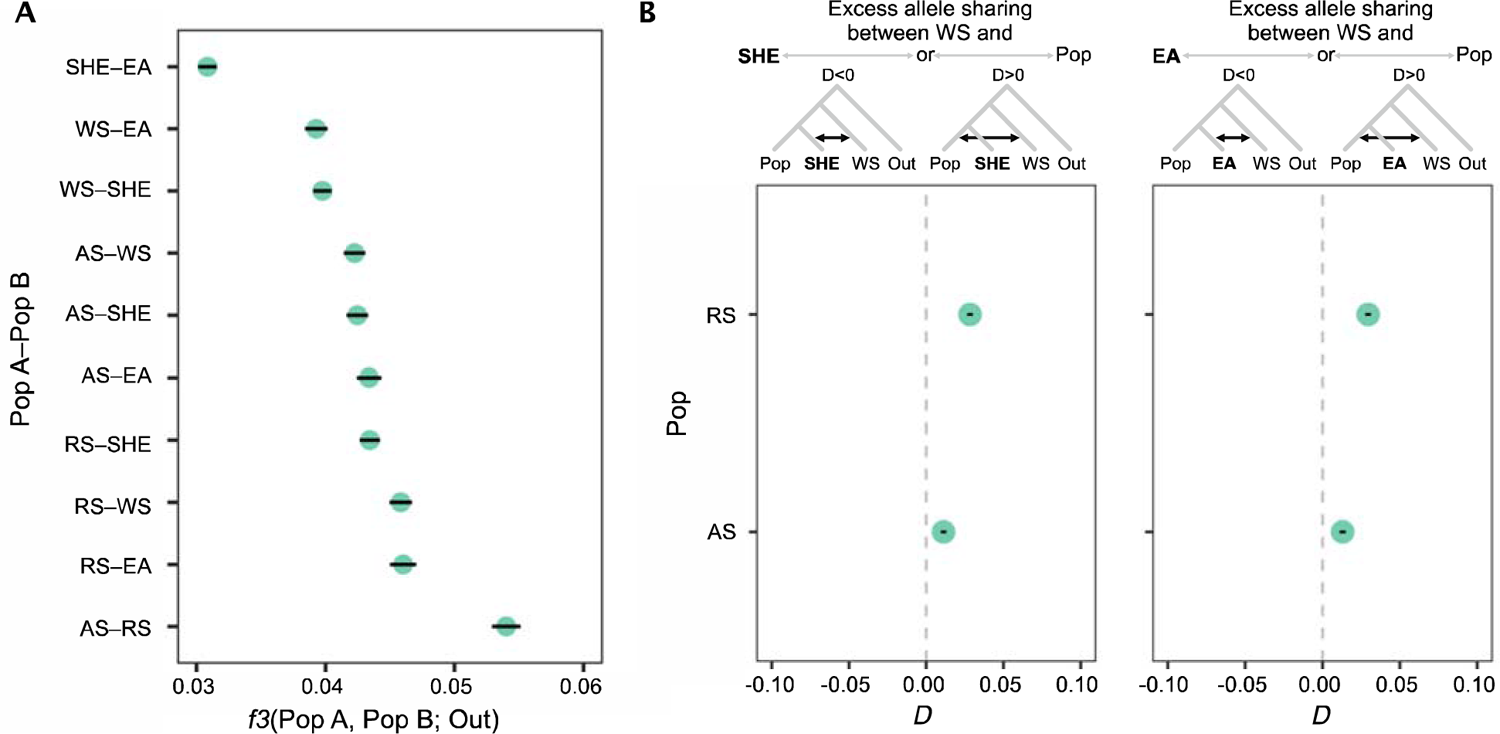
Evidence of distinct allele frequency correlations between Amundsen Sea, Weddell Sea and Ross Sea, as well as contemporary gene flow in *Pareledone turqueti*. Error bars (black horizonal lines) = standard errors, filled circles = significant (Z-score values > 3 or < −3), Out = outgroup population, which includes Shag Rocks and South Georgia (samples combined). Data = 120,857 SNPs dataset. Abbreviations: Weddell Sea (WS), South Shetland Islands (SHE), Amundsen Sea (AS), Ross Sea (RS), East Antarctica (EA). (**A**) Outgroup-ƒ_3_-statistics between pairs of populations. As ƒ_3_ value increases, more derived allele frequency is shared between the pairs of population. (**B**) *D*-statistic (in the form of BABA-ABBA) examines patterns of alleles sharing across four populations, and indicates whether there is excess allele sharing between distinct populations. Left panel: *D*-statistic is presented in the form of *D*(Pop, SHE, WS, Out), which examines whether there is excess allele sharing between SHE and WS (*D*<0; ABBA) or Pop and WS (*D*>0; BABA). Right panel: *D*-statistic is presented in the form of *D*(Pop, EA, WS, Out), which examines whether there is excess allele sharing between EA and WS (*D*<0; ABBA) or Pop and WS (*D*>0; BABA).

A site-frequency-spectrum (SFS)-based, coalescent demographic modelling framework (*fastsimcoal* (32)) was used to test the hypothesis of whether historical trans-Antarctic seaways existed, with populations subsequently influenced by contemporary circumpolar gene flow. For demographic modelling, we included samples from WS, AS, RS and EA with 163,335 SNPs in *P. turqueti*. We employed a hierarchical approach to test for WAIS collapse scenarios while incorporating modern circumpolar gene flow in the models (fig. S5-S6). Step 1 compared contrasting scenarios of past WAIS configurations with circumpolar gene flow following the directionality of the ACC (clockwise). The models incorporated WS, EA, RS and AS experiencing continuous circumpolar gene flow since population divergence. Under these scenarios, after population divergence, WS, AS, RS experienced no, partial, or complete connectivity, followed by modern ACC gene flow linking between WS, EA, RS and AS. Limited differentiation was found between competing scenarios (no, partial, or complete connectivity) at step 1 (fig. S7-S8). Therefore, at step 2, model complexity was increased to model more ecologically realistic scenarios, with circumpolar gene flow following both directionalities of the ACC and ASC (counter-clockwise) for all scenarios (fig. S9-S10). At step 3, unmodelled ancestral size change was further considered to distinguish competing models from the previous step.

The observed SFSs were statistically best explained by the scenario of a complete historical WAIS collapse (Fig. 3A, fig. S11-S12), followed by modern circumpolar gene flow linked to ACC and ASC. The ancestral lineage of WS, AS, RS and EA populations experienced an expansion followed by a bottleneck (Fig. 3A, table S2). During the mid-Pliocene, which ended at ∼3 million years ago (95% CI between 3.6 and 3.5 Mya), the best model supports WS, AS, RS and EA continental shelf locations splitting into four populations with direct asymmetric gene flow detected between WS-AS-RS. This suggests that ancient seaways were likely opened across the WAIS, which directly linked the present day WS, AS and RS, and could only be facilitated by WAIS collapse during past interglacials. The start of the historical, direct WS-AS-RS connectivity began at ∼3.6-3.0 Mya, which supports geological evidence of historical WAIS collapses during the Pliocene (8, 9). The signature of direct WS-AS-RS connectivity ceased between 139 and 54 ka (based on 95% CI; Fig. 3A), and which is in accordance with the end of the Last Interglacial.

**Fig. 3.**
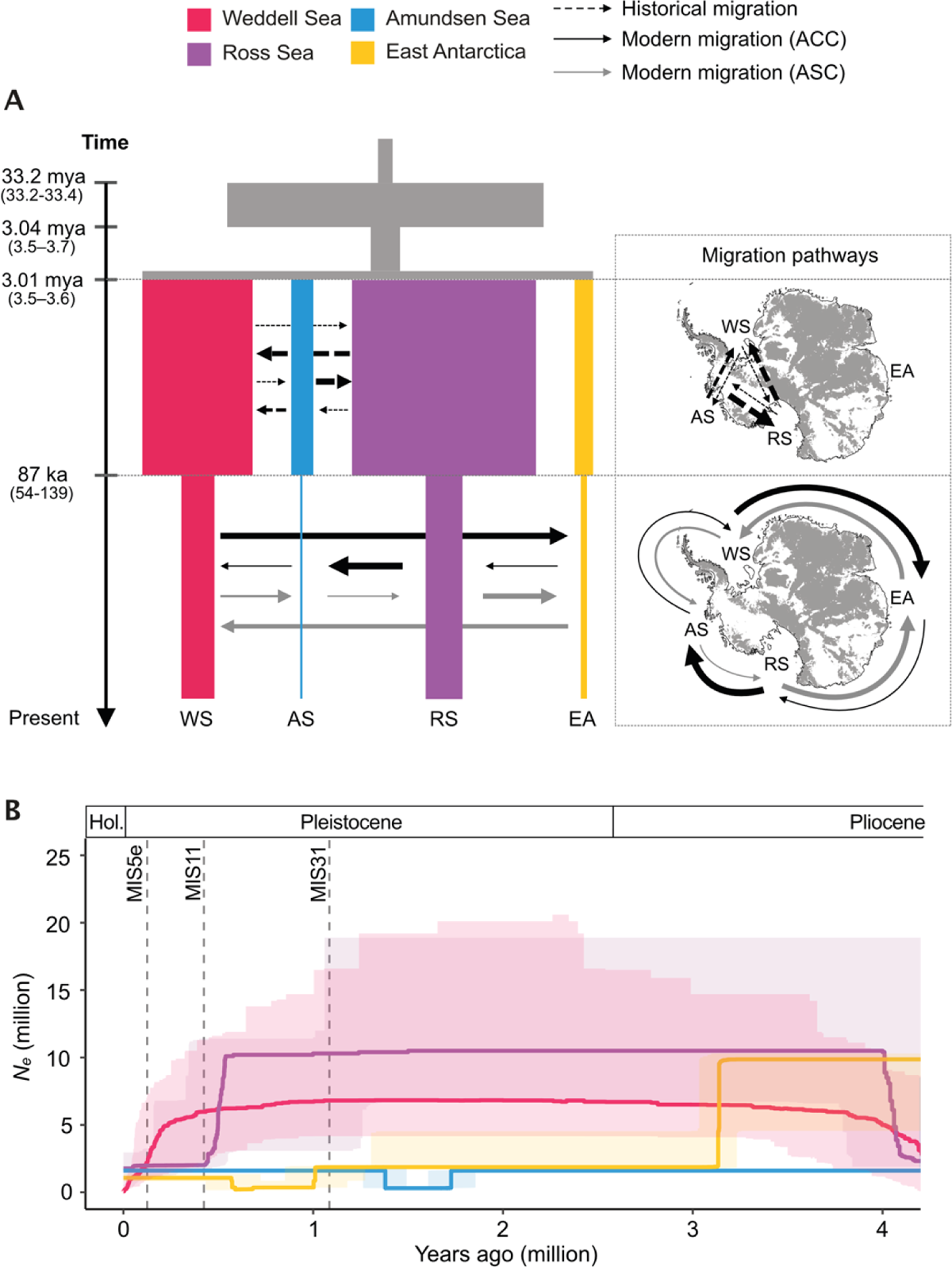
The best-supported demographic model for *Pareledone turqueti* indicated signatures of complete historical West Antarctic Ice Sheet collapse began during the mid-Pliocene until the end of Marine Isotope Stage (MIS) 5e, supplemented by a *StairwayPlot* which indicated past changes in population size. (**A**) Maximum likelihood model for *P. turqueti* including Amundsen Sea (AS), Ross Sea (RS), Weddell Sea (WS) and East Antarctica (EA) populations, shows direct historical gene flow (3 Mya–87 ka) between WS, AS and RS, and modern gene flow (87–0 ka) around the Antarctic continental shelf following the directionality of the Antarctic Circumpolar Current (ACC; clockwise) and Antarctic Slope Current (ASC; counter-clockwise). Maximised parameter estimates are visualised. The associated 95% confidence intervals (CI) are in brackets and reported in Table S2. Time of the events modelled are shown on the left. The width of the bars is proportional to the effective population size of the population. Arrows indicate migration (forward in time), with the width of the arrows proportional to the number of migrants per generation (2Nm). The migration pathways, based on modelled migration directions (forward in time), are also visualised on a map of Antarctica. Map shows sub-glacial bed elevation >0 m above present-day sea level and is extracted from Bedmap2 (33). Data = 163,335 SNPs dataset. (**B**) *StairwayPlot* reconstruction of past changes in effective population size over time in *P. turqueti* in the last ∼4 Mya. Line = median, shaded area = 2.5% and 97.5% confidence limits. Data = 191,024 SNPs dataset. Dashed vertical lines represent timing of Marine Isotope Stage 5e (MIS 5e; ∼125 ka), Marine Isotope Stage 11 (MIS 11; ∼424 ka) and Marine Isotope Stage 31 (MIS 31; ∼1.08 mya). Abbreviation: Holocene, Hol.

Considering the estimated confidence intervals, the cessation of direct gene flow between WS-AS-RS could only be associated to the most recent interglacial MIS 5e (129–116 ka), because the prior super-interglacial (MIS 11; ∼424–395 ka), which would also enable such direct connectivity, is far outside of the upper uncertainty bound of the connectivity. The maximum likelihood value of the direct WS-AS-RS connectivity was dated to 87 ka; the time lag between 87 ka and the time of MIS 5e (129–116 ka) can be explained at least in part by the time it takes for complete trans-West Antarctic migration to influence allele frequencies in a benthic direct developing octopus. Finally, contemporary circumpolar gene flow began after 87 ka, following the directions as the ACC and ASC, which generally reflects the influence of ongoing circumpolar currents on gene flow of Southern Ocean benthic taxa, in particular the ACC (28). For example, the high migration rates observed from WS->EA and RS->AS reflect the strength of ACC relative to ASC. To support the logic that populations were split from the same ancestral population, we considered the alternative tree topologies of diverse 3-population models involving WS, RS and EA or SHE (fig. S13-S14). In all cases, recent, shared ancestry between RS and WS is not an alternative explanation for the observed data (table S3-S4), with consistent outcomes supporting historic direct gene flow between WS-RS ending at around the Last Interglacial. The implication is that the conclusion of WAIS collapse is robust to the model assumption of a split from the same ancestral population.

One of the biggest challenges of inferring demographic events in the late Pleistocene include whether the species experienced a severe bottleneck (i.e. sharp reduction in *N*_e_) in the recent past that eroded genomic history. For example, if the WAIS had collapsed in the late Pleistocene, large areas of newly ice-free habitats (where the WAIS existed previously) would have become available for benthic fauna to disperse and colonise. During the subsequent glacial maximum, as the AIS expanded across the Antarctic continental shelf, the marine shelf habitats would likely be reduced to small, isolated pockets of *in situ* ice-free refugia (34). Such changes in habitat availability would likely lead to severe population bottlenecks and subsequent genetic drift (34). As a result, a recent severe bottleneck could lead to loss of alleles, increased coalescence rate and events only dating back to the time of most recent bottleneck (35) (i.e. Last Glacial Maximum [LGM] at ∼20 ka in *P. turqueti*). Demographic models detected events prior to LGM in *P. turqueti*, suggesting this species did not experience population bottlenecks in the recent past as severe as might be expected for a Southern Ocean benthic species.

We searched for signals of population size fluctuation prior to the LGM in *P. turqueti* using *StairwayPlot* (36), an SFS-based model-free method. We found that the demographic changes dated by *StairwayPlot* generally correspond with the dating of gene flow changes by *fastsimcoal*.

Pronounced demographic changes (observed from Fig. 3B) were detected in the RS and WS populations at ∼435 ka and ∼200 ka. These timings coincide with the glacial stage MIS 12 and the relatively cool interglacial of MIS 7, both of which were subsequently followed by periods of peak warmth during MIS 11 and MIS 5e respectively. In species with population structure, population decline at a particular timepoint in *StairwayPlot* corresponds to signal of demographic change such as changes in population structure and migration rate (37). For the case of *P. turqueti*, the population decline detected in RS around MIS 11 and in WS around MIS 5e corresponds to demographic change likely associated with WAIS collapse. In the *fastsimcoal* model (Fig. 3A), the population decline detected across all populations after the Last interglacial also corresponds to the widely-accepted hypothesis that there would be limited *in situ* ice-free refugia on the Antarctic continental shelf during the LGM, leading to population bottlenecks in benthic species that only survived on the shelf (34) (i.e. the case for *P. turqueti* (23)). These patterns suggest that recent super-interglacials and the LGM likely strongly influenced species demography, particularly in populations associated with the signatures of WAIS collapse.

Estimation of past changes in population size with *StairwayPlot* indicated AS and EA populations experienced relatively stable populations since 3 Mya, while *fastsimcoal* indicated population size changes in these populations; such discordances are likely due to method-specific sensitivity in populations with low sample size in regions that are challenging to collect biological samples (n≤5 in AS and EA). Overall, the timing of demographic events detected by independent inferences also corroborates the timings of major glacial-interglacial fluctuations in the Pliocene and the late Pleistocene, as well as events dated using independent markers (mitochondrial data) (23). Therefore, the dating of WAIS collapse, as seen through the genomic data of *P. turqueti* appears to be robust and unconfounded by noise.

Our demographic modelling approach was specifically designed to test whether trans-West Antarctic seaways existed in the past that could be detected with simple contrasting models. The best-supported demographic model was able to characterize an overview of the historical, direct WS-AS-RS connectivity linked to WAIS collapses through the Pliocene and as late as MIS 5e. While additional periods of seaway closure occurred during other geological periods between 3 Mya and MIS 5e, our analyses are focused on dating the most recent gene flow linked to historical WAIS collapse. The evolutionary history of *P. turqueti* is highly complex and populations would have experienced unique demographic changes associated with each glacial-interglacial cycle throughout the Quaternary. We did not sequentially reconstruct their past changes in population size and connectivity patterns to avoid over-parameterisation. Signatures of recent events are more clearly encoded in allele frequencies. We utilised a target capture approach that can sequence reduced representation genomic data in samples with DNA degradation. Our reduced representation SNP dataset has the power to reflect recent major events dated to around MIS 5e, MIS 11 and the Pliocene. Regardless of the overall challenge of demographic modelling for Southern Ocean species, signatures of a complete WAIS collapse, last detected at around the MIS 5e, were clear in *P. turqueti*.

We provide empirical evidence indicating that the genomic signatures of marine-based sectors WAIS collapse were present during the Last Interglacial (MIS 5e), when GMST was 0.5–1.5 °C warmer than the preindustrial. Future WAIS collapse on centennial timescales is considered as a low likelihood process (4), however, in recent trajectories estimated for temperature rise, such as for the most optimistic emission scenario Shared Socio-economic Pathway (SSP) 1-1.9, the air temperature is projected to reach +1.2–1.7 °C by 2100 (very likely range) (4). Moreover, Antarctic Ice Sheet models simulating the response to Intergovernmental Panel on Climate Change (IPCC) emissions scenarios (15, 38) show a threshold is crossed when warming is above SSP2-2.6 (the Paris climate target), whereby, ice shelves are lost and marine-based sectors of WAIS undergo self-reinforcing melting due to marine ice sheet instability processes. This study provides empirical evidence indicating WAIS collapsed when global mean temperature was similar to today, suggesting the tipping point of future WAIS collapse is close. Future global sea-level rise projections should consider the irreversible collapse of the WAIS, and some marine sectors of the EAIS (39), which will commit the planet to multi-metre GMSL over the coming centuries and millennium if global warming exceeds +1.5–2 °C above preindustrial levels (1, 8, 40).

## Supporting information

Data S1, Data S2, DataS1-S2

## Acknowledgments

We thank the Australian Antarctic Division (AAD), Alfred Wegener Institute for Polar and Marine Research (AWI), British Antarctic Survey (BAS), Museum Victoria (MV), National Institute of Water and Atmospheric Research (NIWA), and G. Jackson for assistance and samples for genetic analysis. We are grateful to T. Jernfors (University of Jyväskylä) for sequencing assistance.

## Funding

Australian Research Council (ARC) Discovery grant DP190101347 (JMS, NGW, NRG, TRN)

New Zealand Ministry of Business, Innovation and Employment through the Antarctic Science Platform (ANTA1801) (NRG, TRN)

Thomas Davies Research grant (Australian Academy of Science) (JMS) David Pearse bequest (SCYL)

Antarctic Science Bursary (SCYL)

Antarctic PhD student support grant (Antarctic Science Foundation) (SCYL)

Australasian eResearch Organisations (AeRO) Cloud Grant (SCYL)

CoSyst grant (JMS, PCW)

Academy of Finland grant 305532 (PCW)

Australian Research Council (ARC) SRIEAS Grant SR200100005 Securing Antarctica’s Environmental Future

Scientific Committee on Antarctic Research (SCAR) INSTANT programme

## Author contributions

Conceptualization: NGW, NRG, TRN, JMS

Methodology: SCYL, NGW, CNSS, JMS

Investigation: SCYL

Formal Analysis: SCYL, IRC

Visualization: SCYL

Funding acquisition: SCYL, NGW, NRG, TRN, PCW, JMS

Resources: PCW, ALA, FCM, KL

Supervision: NGW, CNSS, JMS

Writing – original draft: SCYL

Writing – review & editing: SCYL, NGW, NRG, TRN, PCW, CNSS, IRC, ALA, FCM, KL, JMS

## Data and materials availability

The ddRADseq data of Southern Ocean octopus generated for target capture bait design is deposited on National Centre for Biotechnology Information (NCBI) under the BioProject PRJNA853080, with Sequence Read Archive (SRA) accessions SRR19893055–SRR19893494. The target capture of ddRAD loci data in *Pareledone turqueti* is deposited under the BioProject PRJNA853871, with SRA accessions SRR19892485–SRR19892582. The draft partial genome of *P. turqueti* is available for download from https://www.marine-omics.net/resources/. All software used for data analyses in this study is publicly available. Detailed methods including scripts and command used to perform all analyses are provided at https://github.com/sallycylau/WAIS_turqueti.

## Supplementary Materials

### Materials and Methods

#### Supplementary Text

Figs. S1 to S26

Tables S1 to S13

Data S1 to S2

References (*41*–*96)*

