## Supplementary Materials for "Genomic evidence for West Antarctic Ice Sheet collapse during the Last Interglacial"

**This PDF file includes:**

Materials and Methods

Supplementary Text

Figs. S1 to S26

Tables S1 to S13

References (41–96)

**Other Supplementary Materials for this manuscript include the following:**

Data S1 to S2

Materials and Methods

Target capture sequencing of ddRAD loci in *P. turqueti*

Tissue samples of *Pareledone turqueti* (n = 96) collected around the Antarctic continental shelf and Antarctic islands (Fig. 1A), between the depths of 102–1342 m, were sequenced with target capture probes designed from previously identified ddRADseq loci (see Supplementary Text “Discovery of ddRADseq loci for target capture sequencing”; Data S1-S2). Two outgroup species (*P. aequipapillae*; ID: 44064_1 *and P. cornuta*; ID: CT931) collected from the Ross Sea and Adélie Land, respectively, were also included in the target capture dataset. For target capture sequencing, we did not include the standard DNA shearing step as these samples had already been identified as having degraded DNA. Libraries with unique index adapters were built and pooled into single capture reactions (six libraries per capture). All libraries were enriched in capture reactions using myBaits® following the manufacturer’s protocol and the resulting capture reactions were sequenced on Illumina NovaSeq S4 flow cells with 150 bp paired end reads. Sixty-three *P.* *turqueti* samples examined in this study, as well as the two outgroup samples, were previously included in studies that analysed Cytochrome c oxidase subunit I (COI) and microsatellite data (*23*, *41*).

Target capture data processing, reads mapping and site calling

Raw target capture reads were demultiplexed with adapters and barcodes were removed using *process_shortreads* in *Stacks* v2.3d (*42*). Reads with phred quality less than 20 (Q < 20) were also discarded, and polyG in read tails were trimmed, using *fastp* v0.20 (*43*). Potential contaminants (human and microorganisms) were identified using *Kraken* v1.0 (*44*) and sequences that were classified under the contaminant database (MiniKraken 8GB 2017) were removed. Cleaned and trimmed reads were then checked for quality using *fastQC* v0.11.7 (*45*).

Cleaned target capture reads were mapped to the consensus sequences of ddRAD loci that were used for bait design using *bwa* v0.7 *mem* with default parameters (*46*). *Samtools* v1.7 (*47*) was used to sort alignments by coordinates. PCR duplicates were marked and removed using *picard* v2.18.1 (*48*) to minimise PCR duplicates related errors in downstream analyses. Sites were called across all samples using *bcftools* v1.7 *mpileup* (*49*)*.* Then, indels and samples with high missing data on an individual basis (> 80%) were removed. Further SNP filtering was performed differently based on the assumptions of each type of data analysis (as indicated below) using *VCFtools* v0.1.16 (*50*). The MAPQ (MAPping Quality [Phred-scaled]) score per sample ranged from 20.9 to 35.6, with the exception of one individual sample from South Orkney Islands (ID: PT255) with a low MAPQ score of 13.7. Since samples from South Orkney Islands are hard to access (a total of 2 samples were collected in this study), we retained PT255 and analysed this sample only in genetic diversity statistics, Principal Component Analysis [PCA], *Structure* v.2.3.4 (*51*) and *TreeMix* v.1.13 (*52*) to understand the broad population structure of *P. turqueti*.

For the inference of population structure and relationships at a circumpolar scale (genetic diversity statistics, PCA, *Structure* and *TreeMix*), we included all *P. turqueti* samples (n = 96). We reduced the dataset to 5,188 biallelic unlinked SNPs (hereafter “5,188-SNP dataset”), filtered based on the following steps. Only sites with Phred quality score higher than 30 were kept (--minQ 30). Sites with mean read depth of less than 16x (=average depth (48.2x)/3) and greater than 96x (=2*average depth) across individuals were removed (using --min-meanDP 16 and --max-meanDP 96). Only biallelic sites were kept (--min-alleles 2, --max-alleles 2). Sites were kept if present in at least 50% of all samples (--max-missing 0.5). Sites with a minor allele frequency of at least 5% were kept (--maf 0.05). To strictly remove sites that belonged to paralogous regions and therefore artificial SNPs, sites with a maximum observed heterozygosity of 0.5 were kept (*53*, *54*), identified via the R package *adegenet* v2.1.3 (*55*). Lastly, only one site per locus was kept (--thin 1000; an arbitrary number larger than the longest contig, in basepair [bp] in the bait set).

For the inference of admixture (*AdmixTools* v7.0.1 (*56*)), filtering steps were performed following “5,188-SNP dataset” with variations, resulting in 120,857 biallelic SNPs being kept in the dataset (hereafter “120,857-SNP dataset”). Instead of filtering by minor allele frequency, a minor allele count of 1 was applied to keep variants (--mac 1 in *VCFtools*); and all SNPs per locus were kept.

For demographic modelling using *fastsimcoal* v2.6 (*32*), we reduced the dataset of 163,335 biallelic SNPs based on the following steps (hereafter “163,335-SNP dataset”). Indels and sites with mean read depth of less than 5x and greater than 103x across individuals were removed (--min-meanDP 5, --max-meanDP 103). From here on, only polymorphic sites were kept and filtered based on the following steps. Sites with Phred quality scores higher than 20 (--minQ 20) were kept. Sites with Hardy-Weinberg equilibrium (HWE) threshold of *p* > 0.0001 were kept (--hwe 0.0001). Only biallelic sites are kept (--min-alleles 2, --max-alleles 2). Only sites with a maximum observed heterozygosity of 0.7 were kept, a threshold that removes sites that likely belonged to paralogous regions in RAD data (*53*, *54*), identified via the R package *adegenet*. SNPs were then polarised using outgroup species information (*P. aequipapillae* and *P. cornuta*). Next, we randomly resampled a fixed number of diploid genotypes from each locality (WS: 15, AS: 4, RS: 8, EA: 5) to a polymorphic site dataset without missing data while maximising the number of SNPs and genotypes across localities, using a custom python script fastsimcoal/sampleKgenotypesPerPop.py (http://cmpg.unibe.ch/software/fastsimcoal27/additionalScripts.html). An unfolded multidimensional SFS was generated using the python script fastsimcoal/vcf2sfs.py (https://github.com/marqueda/SFS-scripts/blob/master/vcf2sfs.py). Finally, we manually added the number of monomorphic sites (=1,528,057) to the observed SFS, determined via the total number of sites kept after filtering indels and mean read depths minus the number of polymorphic sites kept after SNP filtering.

For inferring past population size change (*StairwayPlot* v2 (*36*, *57*)), we reduced the dataset to 191,024 SNPs (hereafter “191,024-SNP dataset), without resampling genotypes from each locality following the steps in the “163,335-SNP dataset”. Unfolded 1D-SFS per location was then generated using the python script vcf2sfs.py.

For *AdmixTools*, *fastsimcoal* and *StairwayPlot*, we included *P. turqueti* samples (in diploids) from WS (n = 18), AS (n = 4), RS (n = 10) and EA (n = 5; excluding Adélie Land) (Data S1). Samples from SHE (n = 13), Shag Rocks (n = 9) and South Georgia (n = 19) were also kept for *AdmixTools*. EA localities included Prydz Bay and East Casey Station; we excluded Adélie Land from EA samples as *Structure*, PCA and *TreeMix* indicated uniquely strong admixture between Adélie Land and RS in *P. turqueti*. The observed Adélie Land - RS connectivity could be linked to contemporary regional currents, thus confounding the interpretation of historical trans-West Antarctic connectivity while considering the effects of circumpolar gene flow using EA samples.

Filtering thresholds were adjusted for different datasets in order to retain a maximum number of informative SNPs within the scope of each model’s assumptions while minimizing retention of erroneous loci. For example, hard filtering steps were performed for the “5,188-SNP dataset” with the aim to exclude as many SNPs as possible associated with genotyping errors, linkage and missing data. For the other datasets we retained rare variants (minor allele counts < 2) as they are crucial for the accurate inference of recent demographic events and population size change (*58*). While there is a higher chance of genotyping error with such rare variants the high per-sample sequencing depth in our data ensures that such errors would have had very little overall effect on the allele frequency spectra used for demographic inference.

Genetic structure of *P. turqueti*

To visualise the overall genetic structure of *P. turqueti* at a circumpolar scale, PCA was performed using *adegenet* across all samples. Individual admixture proportions were also estimated via *Structure* (*51*). *Structure* was run between *K* = 1 and 10, with ten replicates per *K* via *Structure_threader* (*59*). Each run was performed with 500,000 iterations and burn-in of 100,000. The meaningful *K* was evaluated based on the highest mean log likelihood [mean LnP(*K*)] and delta*K* statistics using *Structure Harvester* (*60*). *TreeMix* was performed to generate maximum likelihood (ML) tree topology of *P. turqueti*, as well as to model the historical splits and mixture between locations and the amount of genetic drift experienced at each location. For *TreeMix* analysis of *P. turqueti*, outgroup population (samples from South Georgia and Shag Rocks; diverged from other locations since ~4 million years ago (*23*)) were included for tree rooting. *TreeMix* input was generated using the R package *dartR* v1.1.11 (*61*). Within *TreeMix*, migration edges (m) were modelled between 0 and 8, with 10 replicates per m using the bootstrap option with a block size of 1 (assuming loci are unlinked as the input dataset contained one SNP per locus). The optimal number of m was evaluated using Evanno method via the R package *OptM* v0.1.3 (*62*). Among the 10 replicate runs of the best m, the replicate with the highest amount of variance explained was presented. Only the significant migration edges, evaluated using jackknife p values (significance level at 0.05), were presented. Each migration edge was weighted based on the ancestry fraction in the sink population originated from each source population (*52*).

Individual observed heterozygosity (H_o_) was calculated via *dartR*. Genetic differentiation between sample locations was examined with pairwise F_ST_ (Weir and Cockerham [1984] (*63*)) values, calculated using the R package *hierfstat* (*64*). Expected and observed heterozygosity (H_e_, H_o_) and Inbreeding coefficient (F_IS_) across individuals per geographic location were also calculated using *GenoDive* v3.0 (*65*) per location. Locations with low sample size (n ≤ 2 samples; including Antarctic Peninsula, East Casey Station, South Orkney Islands and Bransfield Strait) were omitted from pairwise genetic diversity comparison at a geographical location level.

Test for signatures of isolation-by-water depth

A previous microsatellite study examined the genetic structure of *Pareledone* spp. Suggested the genetic structure of *P. turqueti* could be related to water depth (*41*). We used Generalised Dissimilarity Modelling (GDM) to evaluate whether the genetic structure of *P. turqueti* is associated with water depth, which could confound demographic inferences. GDM is a multivariate statistical method, which uses a nonlinear matrix regression to model the dissimilarity between genetic differentiation and geographic distances and environmental variables (*66*) (hereafter predictors). In GDM, each predictor variable is first transformed using the default three I-spline basis functions, and models are fitted using maximum-likelihood estimation. Variables are standardized, and therefore their resulting coefficients can be compared. The overall model significance and the significance of each predictor was quantified using matrix permutation via the function *gdm.varImp* within the R package *gdm* v1.4.2.2 (*67*). During permutation testing (n = 999), a full model containing all predictors was first considered, followed by iteratively removing the predictor with the lowest coefficient and recalculating the model fit and significance values. Permutation testing is repeated until all non-significant predictors are removed.

Pairwise genetic distance between samples were estimated via *ngsDist* (*68*) using all SNPs across ddRAD loci from the “5,188-SNP dataset” (resulting in 37,322 SNPs, hereafter “37,322-SNP dataset”). For predictor inputs, geographical distances between samples (Euclidean distance; straight direct distance between sample locations) were calculated directly from geographical coordinates within GDM. The environmental variables considered for GDM included sample collection depth. Three samples (ID: JS_40, JS_44, NIWA87470) were omitted from GDM as sample collection depth information was not available. GDM was repeated for samples only from the Scotia Sea (n = 52; including Shag Rocks [n = 9], South Georgia [n = 19], South Orkney Islands [n = 2], Bransfield Strait [n = 1], West Antarctic Peninsula [n = 1], Elephant Island [n = 7], Deception Island [n = 2], King George Island [n = 4], Livingston Island [n = 1], Robert Island [n = 6]) to explore regional signatures of isolation-by-water depth.

Redundancy analysis (RDA) was performed to detect fine-scale genotype-environmental association across samples from the Scotia Sea (n = 52), using the “5,188-SNP dataset”. RDA is a constrained ordination approach that uses multiple linear regression to summarise the linear relationships between genotypes by a set of explanatory environmental predictors. RDA was performed using the R package *vegan* v2.5-6 (*69*). The environmental variables initially considered for RDA included water depth, seafloor water temperature, seafloor water salinity, as well as silicate, phosphate, nitrate and dissolved oxygen (winter and summer values, at the surface and 500 m). Information on water depth, latitude and longitude were obtained from sample information. Values of seafloor water temperature and salinity were calculated from the decadal means of annual average seafloor temperature and salinity at 1^o^ spatial resolution between 1955 and 2010 from World Ocean Atlas 2018 (*70*, *71*). Seafloor temperature and salinity data were estimated based on the information at the depth interval closest to the maximum depth available for each point using QGIS. Values of silicate, phosphate, nitrate and dissolved oxygen (summer and winter values, at the surface and 500 m) were extracted from existing interpolated GIS layers from *Quantarctica* (*72*)*,* gridded at 25km spatial resolution. Multicollinearity between environmental predictors was assessed using the R package *psych* v1.9.12 (*73*), and predictors with low collinearity (r < 0.7) were kept for GDM. After checking for collinearity (r > 0.7), the environmental variables considered included longitude, water depth, seafloor temperature, seafloor salinity and summer nitrate level at 0 m. Results of GDM and RDA are presented and discussed in Supplementary Text “Testing for signatures of isolation-by-water depth”.

Allele frequency correlations between seaway populations

To further explore whether there is direct admixture between seaway populations despite circumpolar gene flow driven by the Antarctic Circumpolar Current (ACC; average speed of 134 Sv [sverdrup] (*74*) and Antarctic Slope Current (ASC; average speed of 10-30 Sv (*29*)), *D*-statistics (in the form of BABA-ABBA) (*30*) and outgroup-ƒ3-statistics (*31*) were calculated using *AdmixTools* (*56*) via *admixr* (*75*). Both tests were constructed based on population trees and were performed between WS, AS and RS populations, with respect to locations (SHE and EA) situated in between WS, AS and RS. Based on mitochondrial data, *P. turqueti* from Shag Rocks and South Georgia are also considered a distinct lineage diverged from Antarctic continental shelf lineage in the mid-Pliocene (*23*). Supporting this concept, *Structure* analysis also indicates samples from this region (Shag Rocks + South Georgia) exhibit genetic isolation. Therefore, for both *D*-statistics and outgroup-ƒ_3_-statistics, samples from Shag Rocks and South Georgia were combined and considered as an outgroup population.

The *D*-statistic examines whether there is excess allele sharing between two of the three ingroup populations, with respect to a common outgroup population. We considered 1) whether there was a partial collapse across WAIS which would result in connectivity between WS and AS, and 2) whether there was also a full collapse across WAIS which would result connectivity between WS and RS. When testing for excess allele sharing between WS and AS or RS, considering SHE or EA, we computed the *D*-statistic of the following form: *D*(seaway population, circumpolar current population, WS, outgroup), where seaway population represented AS or RS, and circumpolar current population represented those would receive potential migration via circumpolar currents (i.e., SHE or EA).

The outgroup-ƒ_3_-statistic examines the branch length between pairs of populations with respect to a common outgroup population. We computed the outgroup-ƒ_3_-statistic of the following form: *f_3_*(Outgroup; A, B), where A and B represented pairs of population between WS, AS, RS, SHE and EA.

For *D*-statistics and outgroup-*f_3_*-statistics, standard errors were computed with block-jackknife procedures, with blocks representing the length of RAD loci. Z-score values > 3 or < -3 were considered significantly different from 0 for both tests. Tabulated outgroup-*f_3_*-statistics and *D*-statistics outputs are presented in table S5-S6.

SFS based inferences – mutation rate and generation time

Event dating in SFS-based inferences is dependent upon generation time and the mutation rate assumed. For site frequency spectrum (SFS)-based inferences (*StairwayPlot*, *fastsimcoal*) for *P. turqueti*, a generation time of 12 years was assumed based on life span (*76*, *77*), as female octopods (including cold water deep-sea octopuses) are known to exhibit a single reproductive period followed by death when eggs hatch (*76*, *78*). The generation time was inferred from beak increments in the most mature *P. turqueti* male collected in Schwarz et al. (*77*). The age of this mature *P. turqueti* specimen was inferred from beak increment counts observed in the closely related *P. charcoti*, which was one growth increment every ~7 days, consistent with previous growth data from a *P. charcoti* individual in captivity (*79*). Anecdotally, a *P. turqueti* specimen was also kept alive for >10 years by Dr Felix Mark, at the Alfred Wegener Institute, Germany, and due to its size was estimated to be older than 2 years when it was first caught. Therefore, the assumption of a generation time of 12 years matches with this observation of *P. turqueti* in captivity. Alternative generation times for *P. turqueti* could include 11 years (based on *P. charcoti*’s age estimation in Schwarz et al. (*76*)) and 13 years (based on the upper limit of *P. turqueti* age estimation in Schwarz et al. (*77*)).

A mutation rate of 2.4 x 10^-9^ per site per generation was used based on the genome-wide mutation rate estimated for the Southern blue-ringed octopus (*Hapalochlaena maculosa*) (*80*).

Demographic modelling

We used demographic modelling to explicitly evaluate whether there were historical migrations linking to no, partial or complete collapse of the WAIS preceding modern-day gene flow in *P. turqueti*. Demographic modelling was performed using the coalescent simulations-based framework in *fastsimcoal*. For demographic modelling, we only considered WS, AS, RS and EA in our models (four-population model), as the model evaluation is based on composite likelihoods which requires a single multidimensional SFS (i.e., four dimensional (4D)-SFS in this study). In a multidimensional SFS with >4 populations, the number of zero entries will increase which makes it challenging for *fastsimcoal* to fit the observed data (*81*). EA samples are chosen to be included in the models instead of SHE as samples from across EA are considered of particular importance in representing clear signatures of circumpolar gene flow (*16*), because they are geographically separated from the WAIS but are also directly influenced by both the ACC and ASC.

Because there are an unlimited number of demographic models to be explored, especially when a high number of populations are incorporated (i.e., four in this study), we used a hypothesis driven, hierarchical approach to reconstruct simple, contrasting demographic models involving no, partial or complete historical collapse of WAIS using *fastsimcoal* (fig. S5). We explored simpler models and subsequently added more complex parameters to improve the fit to the observed data, as recommended by Marchi et al. (*82*), with a hierarchical framework constructed following Marques et al. (*83*). First, we compared six different models comprising different WAIS collapse scenarios (no, partial or complete collapse), while modelling contemporary gene flow driven by the ACC (clockwise) (step 1). These six models had the following conditions: 1) continuous circumpolar gene flow since population divergence (no collapse scenario), 2) strict isolation followed by clockwise circumpolar gene flow (no collapse scenario), 3) gene flow between WS-AS followed by clockwise circumpolar gene flow (partial collapse scenario), 4) gene flow between AS-RS followed by clockwise circumpolar gene flow (partial collapse scenario), 5) gene flow between WS-RS followed by clockwise circumpolar gene flow (full collapse scenario), and 6) gene flow between WS-AS-RS followed by clockwise circumpolar gene flow (full collapse scenario). To infer more ecologically realistic scenarios, we considered complex models (step 2) that included contemporary gene flow following both the directionalities of the ACC (clockwise) and ASC (counter-clockwise). At step 2, we also considered two additional models with the following conditions: 7) gene flow between WS-AS and WS-RS followed by clockwise and counter-clockwise circumpolar gene flow, and 8) gene flow between AS-RS and WS-RS followed by clockwise and counter-clockwise circumpolar gene flow (fig. S5 and S6). Finally, at step 3, we considered an additional ancestral size change among the best competing models from step 2, as *StairwayPlot* indicates population size changes in the ancestral lineage that are unmodelled in step 2 (population expansion and bottleneck) (fig. S15).

Model run and selection

For each model, we performed 100 independent runs of random starting parameter combinations, with each run pooling SFS entries with fewer than 10 SNPs in order to avoid overfitting (-C 10), consisting of 50 ECM optimisation cycles and using 200,000 coalescent simulations. We then re-estimated the likelihoods of each model, based on the maximum-likelihood estimates obtained from the best run (*_maxL.par), also using 100 independent runs and 200,000 coalescent simulations. The re-calculated likelihoods should closely approximate the true likelihoods as they are maximised under each model scenario, and the distribution of the re-calculated likelihoods should reflect the inherent stochasticity of coalescent simulations (*32*). We also introduced upper bound for the parameter T1 (divergence time estimate) to reduce the parameter space within the time period of interest (i.e., history since speciation) in complex models (*83*, *84*). The upper bound of T1 was constrained by the known conservative (median) estimate of species divergence time, which was 4 million years ago for *P. turqueti* (*23*). This divergence estimate was chosen as it was calculated using different markers than RAD loci; these divergence times were estimated using mitochondrial data of *P. turqueti* (*23*).

Model fits were evaluated based on the lowest deltaLikelihood, Akaike's information criterion (AIC) and AIC weights. We also visualised the distributions of re-estimated deltaLikelihood and AIC values in order to assess the variance between model fitting runs. For the final best model, we visually inspected the fit of the observed versus expected SFS, as well as the residuals in model fitting, to evaluate whether the final selected model is approximated to the observed data (see Supplementary Text “Extended results of demographic modelling”).

The 95% confidence intervals (CI) of parameters of the best model were calculated using 100 replicates of non-parametric bootstrapped multiSFS. The 100 bootstrapped replicates were generated via vcf2sfs.py. Within each replicate, the parameters under the best model scenario were estimated with 20 independent runs with 200,000 coalescent simulations, pooling SFS entries with fewer than 10 SNPs (-C 10), and 50 ECM optimisation cycles, with the ML parameter values as starting parameters (-initvalues). The parameter estimates of the best run from each bootstrapped replicate were used to compute the 95% confidence interval with the R package *boot* (*85*).

Is recent shared ancestry between RS-WS an alternative explanation to the data?

In the four-population demographic models, we assume WS, AS, RS and EA are split from the same ancestral population, due to a fundamental geographical constraint; WS and RS are separated by the grounded Antarctic Ice Sheet and occur on opposite sides of West Antarctica. To test this further, we performed additional three-population demographic models between WS, RS and SHE, and between WS, RS and EA, following our assumption in which all populations can or cannot be split from the same ancestral population. Firstly, we explored whether the observed data is best explained by 1) shared recent ancestry between WS and RS, with East Antarctica (EA) or the South Shetland Islands (SHE) as a more divergent population, 2) WS, RS and EA or SHE split from the same ancestral population, or 3) WS, RS and EA or SHE split from the same ancestral population with historical direct connectivity between WS and RS (i.e., full WAIS collapse). Model illustrations are presented in fig. S13.

Then, we explored whether the observed data is also best explained by 1) WS, RS and EA or SHE splitting from the same ancestral population with historical direct connectivity between WS and RS between T1 and T2 (i.e., complete WAIS collapse), followed by present day gene flow, and 2) shared recent ancestry between WS and RS followed by present day gene flow (i.e., no WAIS collapse). Model illustrations are presented in fig. S14. We consider present day gene flow to follow both the ACC (clockwise) and ASC (counter-clockwise).

For demographic modelling between WS, RS and EA, we reduced the dataset to 167,612 biallelic SNPs based on the following steps (hereafter “167,612-SNP dataset”). SNP filtering steps followed the four-population model pipeline until randomly resampling a fixed number of diploid genotypes from each locality (WS: 15, RS: 8, EA: 5) to a dataset without missing data while maximising the number of SNPs and genotypes across localities, using a custom python script fastsimcoal/sampleKgenotypesPerPop.py. An unfolded multidimensional SFS was generated using the python script fastsimcoal/vcf2sfs.py. Finally, we manually added the number of monomorphic sites (=1,519,476) to the observed SFS, determined via the total number of sites kept after filtering indels and mean read depths minus the number of polymorphic sites kept after SNP filtering.

For demographic modelling between WS, RS and SHE, we also reduced the dataset to 195,003 biallelic SNPs based on the following steps (hereafter “195,003-SNP dataset”). SNP filtering steps also followed the four-population model pipeline until randomly resampling a fixed number of diploid genotypes from each locality (WS: 15, RS: 8, SHE: 13) to a dataset without missing data . Unfolded multidimensional SFS was generated using the python script fastsimcoal/vcf2sfs.py. Finally, we manually added the number of monomorphic sites (=1,514,898) to the observed SFS, determined via the total number of sites kept after filtering indels and mean read depths minus the number of polymorphic sites kept after SNP filtering.

Model runs and selection were performed following the steps detailed above for the four-population models. For discussions of these three-population models, see Supplementary Text “Extended results of demographic modelling”.

Analysis of simulated data under the best and worst competing models

Simulated data was generated with *fastsimcoal*, under the best fitted model at step 3 (“anc_psccc_fullcol2” [complete WAIS collapse]), and competing models at step 2 (“psccc_nocol” [no WAIS collapse] and “psccc_parcol” [partial WAIS collapse]) with input parameters set to their maximum likelihood values and mutation rate set to 2.4 x 10^-9^ per site per generation as used in *fastsimcoal* and *Stairwayplot*. SNP data was generated by simulating a large number (100 million) of short (10 bp) fragments from which a subset of 100k SNPs were retained for analysis (-s 100000). The resulting outputs were converted to vcf format using a custom awk script.

To examine whether the simulated data could explain the observed data, we calculated *f*4-statistics of the simulated data and observed data. *f*4-statistics were used for data comparison as we did not model an outgroup population in *fastsimcoal*, which is required for other D-statistics and outgroup-*f*3-statistics. Similar to D-statistics, *f*4-statistics explore the correlations of allele frequency differences between four populations (*56*). *F*4-statistics were computed via *AdmixTools* in the following form: *f*4(W, X; Y, Z), where W, X, Y and Z represent individual populations between WS, AS, RS and EA. Standard errors were computed with block-jackknife procedures, with block size defined as 1 SNP per block. Z-score values > 3 or < -3 were considered significantly different from 0. The results are discussed in Supplementary Text “Analysis of simulated data”.

Past population size changes

Past effective population size (*N*_e_) changes within WS, AS, RS and EA populations of *P. turqueti* were reconstructed using *StairwayPlot*. *StairwayPlot* is a model-flexible method that infers past population size changes over specific points in a genealogy through 1-dimensional SFS (1D-SFS). *StairwayPlot* was chosen to further explore past population size changes instead of demographic models (e.g. *fastsimcoal*) as it is not constrained by a-priori information, which can in turn explore a larger model space than parameterised demographic models (*35*). *StairwayPlot* is also known to reconstruct recent population size changes with high accuracy comparable to whole-genome Sequentially Markovian Coalescent (SMC)-based methods (*86*). The total sequence length was defined as the length of genome explored after SNP filtering (= number of monomorphic sites observed [1,528,057] + number of filtered polymorphic sites kept [191,024]). The percentage of sites used for training was 67% and the number of random break points for each run were (nseq-2)/4, (nseq-2)/2, (nseq-2)*3/4, nseq-2 based on default values. Each run was performed with a random starting seed.

To evaluate whether the constant population size observed in AS and EA in Fig. 3B is due to low sample size, samples of WS and RS were downsampled to the same sample size as AS (2N = 4) and EA (2N = 5) via sampleKgenotypesPerPop.py. *StairwayPlot* was then performed on the downsampled WS and RS datasets following the above steps. See supplementary text “Effect of sample size in *StairwayPlot*” for extended results.

Supplementary Text

Discovery of ddRADseq loci for target capture sequencing

*Draft reference genome sequencing and assembly*

A draft reference genome of *P. turqueti* was sequenced from two individuals collected from Elephant Island (ID: PT186) and the South Orkney Islands (ID: PT244) (Data S2). Total genomic DNA of both of these samples (gDNA) was extracted using a Dneasy Blood and Tissue Kit (Qiagen), following the manufacturer’s protocol. Sample PT186 was sequenced on PacBio Sequel system (20 K insert library) with three cells which generated a total read volume of 28 Gigabase pairs (Gbp). Sample PT244 was sequenced using both 200 base pair (bp) and 500 bp insert libraries on an Illumina HiSeq X ten in 150 bp paired-end mode. One flow cell was used for the 200 bp library and two flow cells for the 500 bp. Genome size was estimated at between 3.7 Gb and 8.1 Gb based on the Illumina reads using Genomescope 2.0 (*87*). Genome assembly was performed with *Flye* v2.4 (*88*) using the long-reads from PT186 and then error corrected using reads from PT244 with *Pilon* (*89*). The final assembly had a total length of 517 Mb from 38,290 contigs with the largest contig of 146 Kb and N50 of 16.9 Kb. It is available for download from https://www.marine-omics.net/resources/ (Direct download link from host: https://cloudstor.aarnet.edu.au/plus/s/opg7MQ0tVHtCmMN/download). We found that the mapping rate of Illumina raw reads from sample PT244 to the polished assembly was high (~92%) indicating that despite the small size of this assembly it captured the vast majority of unique genomic sequence. Nevertheless, the small assembly size compared with estimated genome size suggests that the assembly is highly incomplete, probably due to the collapse of many repetitive regions. We therefore used it purely for the purpose of identifying ddRAD loci for target capture sequencing.

### *ddRAD library preparation, sequencing and SNP calling*

As part of a wider effort to perform ddRAD sequencing across different Southern Ocean octopus species, 440 Southern Ocean octopus specimens (*Adelieledone polymorpha, A.* *adelieana*, *Adelieledone* sp., *Pareledone turqueti*, *P. aequipapillae*, *P. prydzensis*, *P. cornuta*, *P. subtilis*, *Pareledone* sp., *Megaleledone setebos* and *Graneledone* sp.) (Data S2) were selected for ddRADseq library preparation and sequencing. ddRADseq libraries were prepared at the Beijing Genomics Institute (BGI) Tech Solutions Co. Limited (Hong Kong) following Peterson et al. (*25*). Briefly, genomic DNA of each sample was digested with MseI and EcoRI restriction enzymes, ligated with barcoded adapters, pooled digested ligated fragments were size selected using Blue Pippin and divided into libraries. Twenty-two technical replicates were also included across libraries (see Data S2). All libraries were amplified via PCR using indexed primers and sequenced on a HiSeq X ten at BGI.

Raw ddRAD reads were demultiplexed with barcodes and adapters removed by BGI using their in-house pipeline. Reads with phred quality less than 20 (Q < 20) were also discarded using fastp. Potential contaminants (human and microorganisms) were identified using *Kraken*, and reads that matched those of the contaminant database were removed. Cleaned and trimmed reads were checked for quality using *fastQC*, and mapped to the draft genome of *P. turqueti* using *bowtie2* v2.3.4.1 (*90*) (--very-sensitive-local). Local alignment (--very-sensitive-local) was used, following Souza et al. (*91*), since the ddRADseq dataset contains a wide variety of Southern Ocean octopod taxa that may contain structural rearrangements or variants at either ends of reads that are different from the reference (*P. turqueti*). *Samtools* was used to sort the alignments (BAM files) by coordinates. ddRAD loci were built from aligned and sorted reads, and SNPs were called, using the *Stacks* v2.3d *gstacks* module with default settings (*92*).

### *ddRAD loci discovery for target capture sequencing of* P. turqueti

Initial assessment of raw genotype calls from *Stacks* indicated 155 out of 440 Southern Ocean octopus samples exhibited a high amount of missing data (> 80%), with 92 out of these 155 samples identified as *P. turqueti*. Samples with high levels of missing data were likely degraded due to long term storage. Then, a target capture bait set was designed with the intention of capturing a high proportion of the same loci in the degraded samples that were included in the ddRADseq (non-degraded) dataset. Loci discovery for this purpose was performed using a total of 285 samples comprising those with missing data less than 80% and included samples from the following: *A. adelieana* (n = 4), *A. polymorpha* (n = 1), *Adelieledone* sp*.* (n = 12), *P. turqueti* (n = 204), *P. aequipapillae* (n = 28), *Pareledone* sp. (n = 15), *M. setebos* (n = 3), *Graneledone* sp (n = 1), as well as technical replicates (n = 17). (Data S2). The Stacks population module was then performed to retain sites that were present in 50% of the remaining samples (-R 0.5) with at least a minor allele frequency of 0.01 (-min-maf 0.01), which resulted in 31,142 loci retained. Discriminant analysis of principal components (DAPC) was performed via the R package *adegenet* to visualise potential batch effects between libraries (no batch effect was found). When the technical replicates were paired together, the replicate with the highest amount of missing data was removed.

The consensus fasta sequences of the 31,142 loci were then aligned back to the reference *P. turqueti* genome using *bowtie2* with end-to-end alignment (--sensitive). Of the 31,142 loci, 8,942 loci were aligned back to the genome exactly once, while 20,123 loci aligned multiple times. Only the 8,942 uniquely aligned loci were retained for target capture bait design, to avoid paralogous regions which can compromise phylogenetic inference (*93*).

### *Bait design for the target capture sequencing of ddRAD loci in degraded* P. turqueti *samples*

The consensus sequences of the filtered ddRAD loci (n = 8,942) were used for custom biotinylated RNA bait manufacturing at Arbor Bioscience (Ann Arbor, MI, USA). Input sequences were soft-masked (0.5%) for simple repeats and low-complexity regions using *Repeat Masker* (*94*), and candidate bait sequences were designed based on bait length (70 nucleotides per bait) and 3 X tiling per locus. Candidate baits were removed if, 1) they were greater than 25% soft-masked for simple repeats, 2) had hits to regions of the *P. turqueti* genome (this study) and the common octopus *Octopus vulgaris* genome (GenBank assembly accession: GCA_003957725.1) (*95*) that were greater than 25% soft-masked (i.e., repeats and low-complexity regions), or 3) failed the Arbor Bioscience in-house moderate Basic Local Alignment Search Tool (BLAST) parameters, which take into account the BLAST hit for a bait and predicted melting temperatures. The *Octopus vulgaris* genome was also included when searching for repeats and low-complexity regions as 1) this is the only available complete octopus genome to date closest to our study species, as well as to 2) ensure octopod-related simple repeats and low-complexity regions were removed given we recognise our draft genome is incomplete. The final myBaits© (Arbor Bioscience) panel contained 86,422 baits that targeted 8,877 ddRAD loci with at least one bait.

Extended results of demographic modelling

*Demographic modelling of* P. turqueti

We used a hierarchical approach to build a demographic model of Weddell Sea (WS), Amundsen Sea (AS), Ross Sea (RS) and East Antarctica (EA) populations in *P. turqueti*. We started from simple models (step 1; models only including contemporary gene flow flowing in the direction of the Antarctic Circumpolar Current [ACC]) and then increased model complexity (step 2; models including contemporary gene flow flowing in the direction of the ACC and the Antarctic Slope Current [ASC], and step 3; models with ancestral size changes) (fig. S5 and S6). At step 1, a limited and overlapping differentiation between maximised Akaike information criterion (AIC) values (median between 2528639 and 2528949) was observed across models (table S7, fig. S7 and S8). This indicates no best model could be distinguished at step 1. Therefore, we further evaluated step 2 models to increase complexity and to model more ecologically realistic scenarios. After incorporating competing scenarios of historical WAIS configurations and contemporary gene flow following the ACC and ASC, the “psccc_fullcol2”, “psccc_nocol” and “psccc_parcol” model were identified as the best competing models (table S8, fig. S9 and S10). At step 3, we included an additional ancestral size change to consider for unmodelled event in the ancestral lineage, which would help to distinguish the competing isolation with migration-based models (“anc_psccc_fullcol2”, “anc_psccc_parcol”) versus secondary contact-based models (“anc_psccc_nocol”) (*96*). The “anc_psccc_fullcol2” model was found to be the best fitted model to the observed data (Table S9, fig. S11 and 12).

We designed simple, contrasting models to specifically test whether trans-West Antarctic seaways existed in the past, and did not model for sequential demographic changes associated with each glacial-interglacial cycle throughout the Quaternary. Overall, we obtained a reasonable fit of the expected and the observed site-frequency-spectrum (SFS) for *P. turqueti* under the step 3 model “anc_psccc_fullcol2” model (fig. S16). Among the SFS entries (fig. S17), there is a good fit of the expected SFS for the entries with more SNPs, with the fit of the expected SFS gradually getting poorer for entries with fewer SNPs. The poorest fits of the expected SFS were observed for the entries with a high number of derived alleles in some populations (fig. S16). This is expected as the modelled demographic scenarios aim to test simple contrasting hypothesised scenarios of whether there was no, partial or complete historical Western Antarctic Ice Sheet (WAIS) collapse, as well as accounting for the parameters of circumpolar gene flow, across four populations (WS, AS, RS and EA). We did not model for detailed demographic changes for each population in order to avoid over-parameterising the models. The unmodelled high number of derived alleles in some populations likely represent unmodelled population-level changes throughout the Quaternary glacial-interglacial cycles.

Exploring the upper and lower limits of estimated generation times (11 and 13 years) did not greatly change the gene flow timing in the best WAIS collapse “anc_psccc_fullcol2” model. For example, the estimation for T3 (time when the signatures of WAIS collapse ceases, when contemporary gene flow between WS-AS-RS-EA linked to circumpolar currents begins) ranges between 80-94 ka based on lower (11 years) and upper estimate (13 years) of generation times, which is similar to the time (=87 ka) estimated using a generation time of 12 years (table S10).

*Evaluation of simpler step 2 models with no or partial WAIS collapse, relative to the best complex model with complete WAIS collapse at step 3*

Evaluation of models at step 2 without complete trans-west Antarctic seaway migration (i.e., no WAIS collapse “psccc_nocol” and partial WAIS collapse “psccc_parcol”) indicate these models were a poorer fit to the observed data based on (fig. S18 to 21), compared to the best and more complex step 3 model with complete WAIS collapse “anc_psccc_fullcol2” (fig. S22). By comparing the 30 worst fitted SFS entries across models, the best and more complex step 3 model with full WAIS collapse (“anc_psccc_fullcol2”) has smaller differences between the observed and expected likelihoods per SFS entry, smaller count differences between the observed and expected likelihoods per SFS entire and a better relative fit between observed and expected entries (fig. S22). Therefore, the best and more complex step 3 model with full WAIS collapse (“anc_psccc_fullcol2”) is a better fitted model.

*Recent shared ancestry is not an alternative explanation of gene flow between Weddell Sea and Ross Sea* – 3 population models

Given our best four-population model results (Fig. 3) suggest direct historical connectivity between Weddell Sea (WS) and Ross Sea (RS), we performed additional three-population models to be certain that the signatures of direct gene flow between WS and RS could not be explained by recent shared ancestry.

When exploring the relationship between RS, WS and EA without present-day gene flow (fig. S13), *fastsimcoal* indicated the scenario of a full WAIS collapse linking gene flow between WS and RS (i.e., model WS_RS_EA_fullcol1) was better supported relative to the scenario of no collapse (i.e., model WS_RS_EA with distinct lineages and EA_WSRS with shared ancestry considered) (table S3). Similarly, when exploring the relationship between RS, WS and SHE without present-day gene flow (fig. S13), *fastsimcoal* also indicated the scenarios of a complete WAIS collapse linking gene flow between WS and RS (i.e., WS_RS_SHE_fullcol1) and WS, RS and SHE as distinct lineages (i.e., SHE_WSRS) were better supported, relative to the scenario of no collapse with shared ancestry between RS and WS considered (i.e., WS_RS_SHE) (table S3). Limited differentiation was detected between WS_RS_SHE_fullcol1 and SHE_WSRS.

Then, we explored whether the observed data is also best explained by 1) WS, RS and EA or SHE splitting from the same ancestral population with historical direct connectivity between WS and RS between T1 and T2 (i.e., complete WAIS collapse), followed by present day gene flow, and 2) shared recent ancestry between WS and RS followed by present day gene flow (i.e., no WAIS collapse) (fig. S14).

When exploring the relationship between RS, WS and EA with present day gene flow (fig. S14), *fastsimcoal* indicated the complete WAIS collapse model (with WS, RS and EA split from the same ancestral population, historical direct connectivity between WS and RS, followed by present day gene flow) was better supported, relative to the model of no collapse with shared ancestry between WS-RS followed by present day gene flow (table S4). Similarly, when exploring the relationship between RS, WS and SHE with present day gene flow (fig. S14), *fastsimcoal* also indicated the complete WAIS collapse model (with WS, RS and SHE split from the same ancestral population, historical direct connectivity between WS and RS, followed by present day gene flow) was better supported, relative to the model of no collapse with shared ancestry between WS-RS followed by present day gene flow (table S4).

Finally, the maximum likelihood parameters from the three-population full WAIS collapse model considering EA (WS_RS_EA_fullcol1_cc) indicated direct connectivity between RS and WS from 3.9 million years ago until 115 ka (around the Last Interglacial), followed by present day gene flow until now (table S11). The maximum likelihood parameters from the three-population full WAIS collapse model considering SHE (WS_RS_EA_fullcol1_cc) indicated direct connectivity between RS and WS from 3.9 million years ago until 282 ka (around the Last Interglacial), followed by present day gene flow until now (table S12).

Overall, all three-population models established that the direct historical gene flow between RS-WS cannot be alternatively explained by recent shared ancestry between RS-WS followed by contemporary circumpolar gene flow around the West Antarctic (represented by SHE) and the East Antarctic (represented by EA) coastline.

*Evaluation of simulated data under the best and competing models*

Using simulated data generated under the best fit *fastsimcoal* model indicating step 3 full WAIS collapse (“anc_psccc_fullcol2”), and the competing step 2 model with partial (“psccc_parcol”) and no (“psccc_nocol”) WAIS collapse, the *f*4-statistics indicate the best model (“anc_psccc_fullcol2”) can explain the observed data better (fig. S23). For *f*4(WS, RS, AS, EA) results of competing models are slightly closer to the observed data, but for *f*4(WS, AS, RS, EA) and *f*4(WS, EA, AS, RS) the full WAIS collapse model gave a result much closer to the observed data than other models. In the case of *f*4(WA, EA, AS, RS) the D values obtained from real data and from the full WAIS collapse model were identically near 0 and non-significant.

Testing for signatures of isolation-by-water depth

The results of GDM show sample collection depth is not a significant factor in explaining the genetic differentiation of *P. turqueti* at a circumpolar level and across the Scotia Sea (table S13). Within the Scotia Sea, we find that samples from Robert Island (n = 6) from South Shetland Islands (SHE) group exhibit a strong association with water depth (fig. S24). Additionally, Structure also shows Robert Island samples exhibit distinct substructure (fig. S25). It is possible the association with water depth could be due to the area’s complex topography and oceanic currents as speculated in Strugnell et al. (2017). To test this possibility, we performed simpler 3 population demographic models substituting SHE and EA in relation to WS and RS (fig. S13-14). We found the same results for both analyses, which again supports the conclusion of the four-population model in the main text (Fig. 3), i.e., signature of full WAIS collapse ceased around the Last Interglacial.

Effect of sample size in *StairwayPlot*

We found that the downsampled WS and RS datasets (to 2N = 5 and = 4) exhibit population sizes near constant, particularly in the recent times (fig. S26). This indicates the *StairwayPlot* inference of the observed AS and EA data (Fig. 3, fig. S15) is likely impacted by small sample size.

 **Fig. S1.**

**Principal component analysis (PCA) of *Pareledone turqueti*.** Samples are separated by geographical locations showing the genetic variation on the first two PC axes (5,188-SNP dataset).

Fig. S2.

***TreeMix* maximum likelihood (ML) tree of *Pareledone turqueti* rooted with outgroup population (Shag Rocks and South Georgia).** Horizontal branch lengths are proportional to the amount of genetic drift that has occurred on each branch. Migration edge is coloured based on migration weight, which corresponds to the % ancestry in the sink population originated from the source population. Only the edges found to be significant by jackknife significance tests were presented. Data = 5,188-SNP dataset. (**A**) ML tree of *P. turqueti*. Terminal nodes are subdivided based on distinct geographical locations. (**B**) Residual matrix visualising the fit of the *TreeMix* modelled allele frequencies to the observed allele frequencies. Residuals are shown as the standard error (SE) of the covariance deviation. Positive residuals (> 0) represent that the *TreeMix* model underestimated the observed covariance, and that the paired populations are more closely related than modelled. Negative residuals (< 0) represent that the *TreeMix* model overestimated the observed covariance, and that the paired populations are more distant than modelled. However, negative residuals are also products of positive residuals being present in the matrix. This topology explains 77.6% of total genetic variance. High SE is observed in locations (Bransfield Strait, South Orkney Islands, East Casey Station, West Antarctic Peninsula) with n ≤ 2.

**Fig. S3.** **Boxplot representing the range of individual observed heterozygosity (H_o_) per sampled locations in *Pareledone turqueti*.** Box represents first and third quartiles. Line within box represents the median. Upper whisker extends to the largest values no more than 1.5*inter-quartile range, and lower whisker extends to the smallest values no more than 1.5*inter-quartile range. Points beyond whiskers are outlying points. Data = 5,188-SNP dataset.

**Fig. S4.**  **Heatmap representing pairwise F_ST_ (Weir and Cockerham) comparisons between sampled locations of *Pareledone turqueti*.** Map of sample locations (examined for pairwise F_ST_) are presented in the left panel. Heatmap representing pairwise F_ST_ comparisons between locations are presented in the right panel. Data = 5,188-SNP dataset.

Fig. S5.

**Hierarchical demographic modelling approach to deduce signatures of historical trans-West Antarctic seaways connectivity in *Pareledone turqueti*.** Simple, contrasting scenarios of past West Antarctic Ice Sheet (WAIS) configurations were compared. (**A**) For the models at step 1, it was hypothesised that since population divergence WS, AS, RS may have experienced any, partial, or complete connectivity, followed by modern circumpolar gene flow linking WS, AS, RS and EA. The possibilities of population size change over each time interval were also considered. For circumpolar gene flow, simpler models which only consisted of the directionality of the Antarctic Circumpolar Current (ACC; clockwise flowing) were performed. (**B**) To increase model complexity, step 2 models were further performed, which considered more complex models that included both directionalities of the ACC and Antarctic Slope Current (ASC; counter-clockwise flowing). (**C**) At step 3, an additional size change in the ancestral population was performed to distinguish between the best competing models at step 2; which include the isolation with migration based models (“anc_psccc_fullcol2”, “anc_psccc_parcol”) versus secondary contact based models (“anc_psccc_nocol”). Each model is labelled by the text above it. Text within each model denotes the parameter labels associated with the population size change at a particular interval (Nxxx), as well as the timing of modelled events (Tx). Dashed lines represent a distinct time interval. Arrows represent migration between populations.

**Fig. S6.**

**Illustrations of the contrasting four-population demographic models to deduce signatures of historical trans-West Antarctic seaways connectivity in *Pareledone turqueti*.** (**A**) At step 1 of the hierarchical demographic modelling approach, contrasting scenarios of past Western Antarctic Ice Sheet (WAIS) configurations were compared. It was hypothesised that since population divergence, the WS, AS and RS may have experienced any, partial, or complete connectivity (black arrows), followed by contemporary circumpolar gene flow driven by the Antarctic Circumpolar Current (ACC, clockwise flowing; grey thick arrows) linking between the WS, AS, RS and EA. (**B**) At step 2, more complex models were further considered; they included both directionalities of the circumpolar gene flow, driven by the ACC (grey thick arrows) and Antarctic Slope Current (ASC, counter-clockwise flowing; grey thin arrows). Each model is labelled by the text above it. Maps illustrate ice thickness of the modern Antarctic Ice Sheet and are extracted from Bedmap2 (*40*).

**Fig. S7**.

**Comparisons of demographic models at step 1 in *Pareledone turqueti*.** See fig. S3 for visualisations of the models. The distributions of loglikelihood (lhood) from 100 independent *fastsimcoal* run (violin plot), with each approximated using 200,000 coalescent simulations under the parameters that maximised the likelihood for each model. Each box represents the interquartile range (25^th^ and 75^th^ percentile), each line represents the median, each dot represents outlier values > 1.5x and < 3x the interquartile range. Lower likelihoods (i.e., better likelihoods) are closer to the origin of the axis. Data = 163,335-SNP dataset.

**Fig. S8.**

**Comparisons of demographic models at step 1 in *Pareledone turqueti*.** See fig. S3 for visualisations of the models. The distributions of loglikelihood (lhood) from 100 independent *fastsimcoal* run (violin plot), with each approximated using 200,000 coalescent simulations under the parameters that maximised the likelihood for each model. Each box represents the interquartile range (25^th^ and 75^th^ percentile), each line represents the median, each dot represents outlier values > 1.5x and < 3x the interquartile range. Lower AIC (i.e., better AIC) are closer to the origin of the axis. Data = 163,335-SNP dataset.

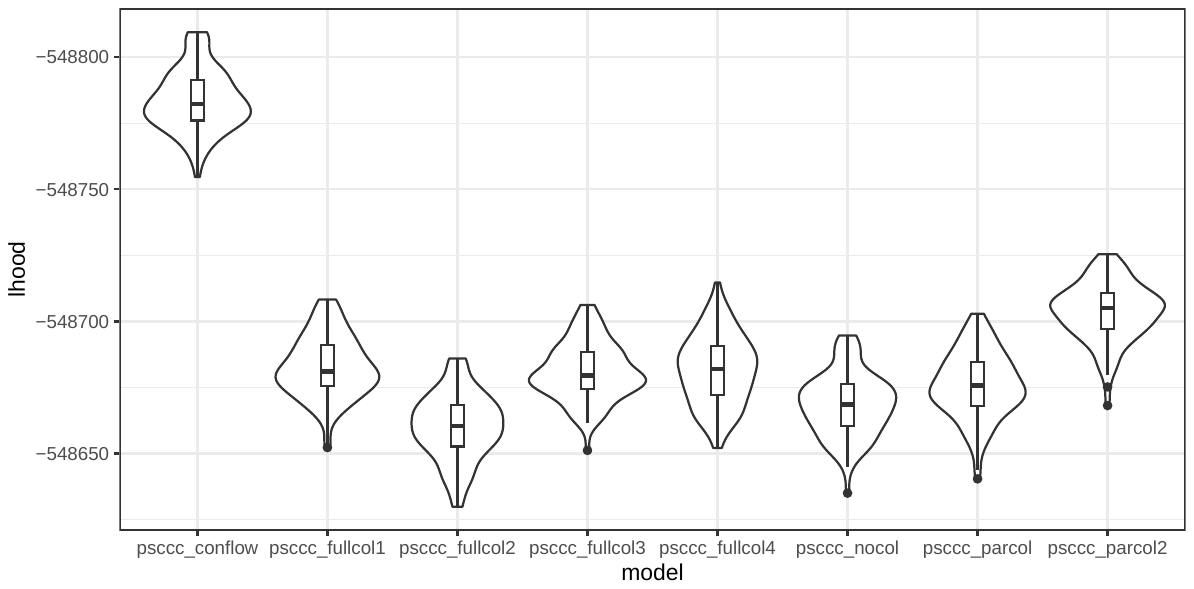

**Fig. S9.**

**Comparisons of demographic models at step 2 in *Pareledone turqueti*.** See fig. S3 for visualisations of the models. The distributions of loglikelihood (lhood) from 100 independent *fastsimcoal* run (violin plot), with each approximated using 200,000 coalescent simulations under the parameters that maximised the likelihood for each model. Each box represents the interquartile range (25^th^ and 75^th^ percentile), each line represents the median, each dot represents outlier values > 1.5x and < 3x the interquartile range. Lower likelihoods (i.e., better likelihoods) are closer to the origi^n^ of the ^ax^is. Data = 163,335-SNP dataset.

**Fig. S10.**

**Comparisons of demographic models at step 2 in *Pareledone turqueti*.** See fig. S3 for visualisations of the models. The distributions of AIC from 100 independent *fastsimcoal* run (violin plot), with each approximated using 200,000 coalescent simulations under the parameters that maximised the likelihood for each model. Each box represents the interquartile range (25^th^ and 75^th^ percentile), each line represents the median, each dot represents outlier values > 1.5x and < 3x the interquartile range. Lower AIC (i.e., better AIC) are closer to the origin of the axis. Data = 163,335-SNP dataset.

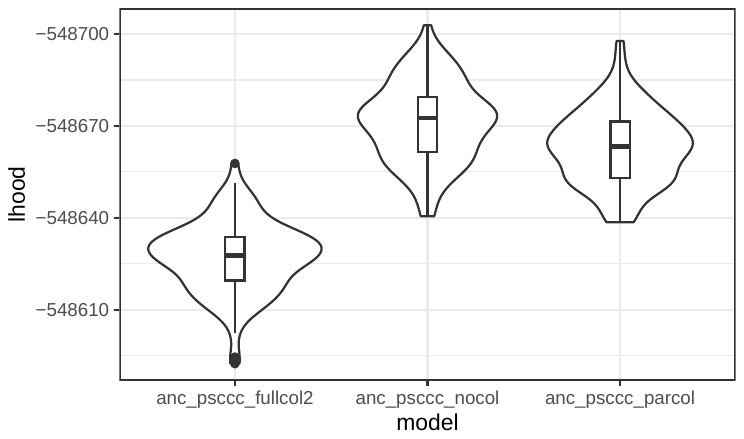

**Fig. S11.**

**Comparisons of demographic models at step 3 in *Pareledone turqueti*.** See fig. S3 for visualisations of the models. The distributions of loglikelihood (lhood) from 100 independent *fastsimcoal* run (violin plot), with each approximated using 200,000 coalescent simulations under the parameters that maximised the likelihood for each model. Each box represents the interquartile range (25^th^ and 75^th^ percentile), each line represents the median, each dot represents outlier values > 1.5x and < 3x the interquartile range. Lower likelihoods (i.e., better likelihoods) are closer to the origi^n^ of the ^ax^is. Data = 163,335-SNP dataset.

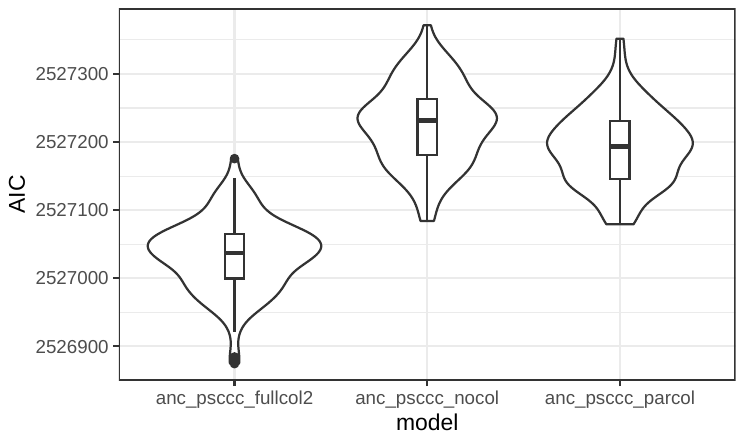

**Fig. S12.**

**Comparisons of demographic models at step 3 in *Pareledone turqueti*.** See fig. S3 for visualisation of the models. The distributions of AIC from 100 independent *fastsimcoal* run (violin plot), with each approximated using 200,000 coalescent simulations under the parameters that maximised the likelihood for each model. Each box represents the interquartile range (25^th^ and 75^th^ percentile), each line represents the median, each dot represents outlier values > 1.5x and < 3x the interquartile range. Lower AIC (i.e., better AIC) are closer to the origin of the axis. Data = 163,335-SNP dataset.

**Fig. S13.**

**Simple models considering for three populations with alternative tree topologies.** (**A**) RS, WS and EA. (**B**) RS, WS and SHE. Models were designed to distinguish whether the conclusion of past gene flow between RS and WS (Fig. 3 in Main text) is robust to the assumption of a split from the same ancestral population. Specifically, contrasting models examined if the observed data is best explained by 1) shared recent ancestry between WS and RS, with EA/SHE as a distant population, 2) WS, RS and EA/SHE split from the same ancestral population, or 3) WS, RS and EA/SHE split from the same ancestral population with historical direct connectivity between WS and RS between T1 and T2. Abbreviation: Ross Sea (RS), Weddell Sea (WS), East Antarctica (EA), South Shetland Islands (SHE). Each model is labelled by the text above it. Text within each model denotes the parameter labels associated with the population size change at a particular interval (Nxxx), as well as the timing of modelled events (Tx). Dashed lines represent a distinct time interval. Arrows represent migration between population.

**Fig. S14.
Simple models considering for three populations with alternative tree topologies, with present day gene flow driven by the Antarctic circumpolar current (clockwise) and Antarctic slope current (counter-clockwise).** (**A**) RS, WS and EA. (**B**) RS, WS and SHE. Models were designed to distinguish whether the conclusion of past gene flow between RS and WS (Fig. 3 in Main text) is robust to the assumption of a split from the same ancestral population. Specifically, contrasting models examined if the observed data is best explained by 1) WS, RS and EA/SHE split from the same ancestral population with historical direct connectivity between WS and RS between T1 and T2 (i.e., full WAIS collapse), followed by present day gene flow, and 2) shared recent ancestry between WS and RS followed by present day gene flow (i.e., no WAIS collapse). Abbreviation: Ross Sea (RS), Weddell Sea (WS), East Antarctica (EA), South Shetland Islands (SHE). Each model is labelled by the text above it. Text within each model denotes the parameter labels associated with the population size change at a particular interval (Nxxx), as well as the timing of modelled events (Tx). Dashed lines represent a distinct time interval. Arrows represent migration between populations.

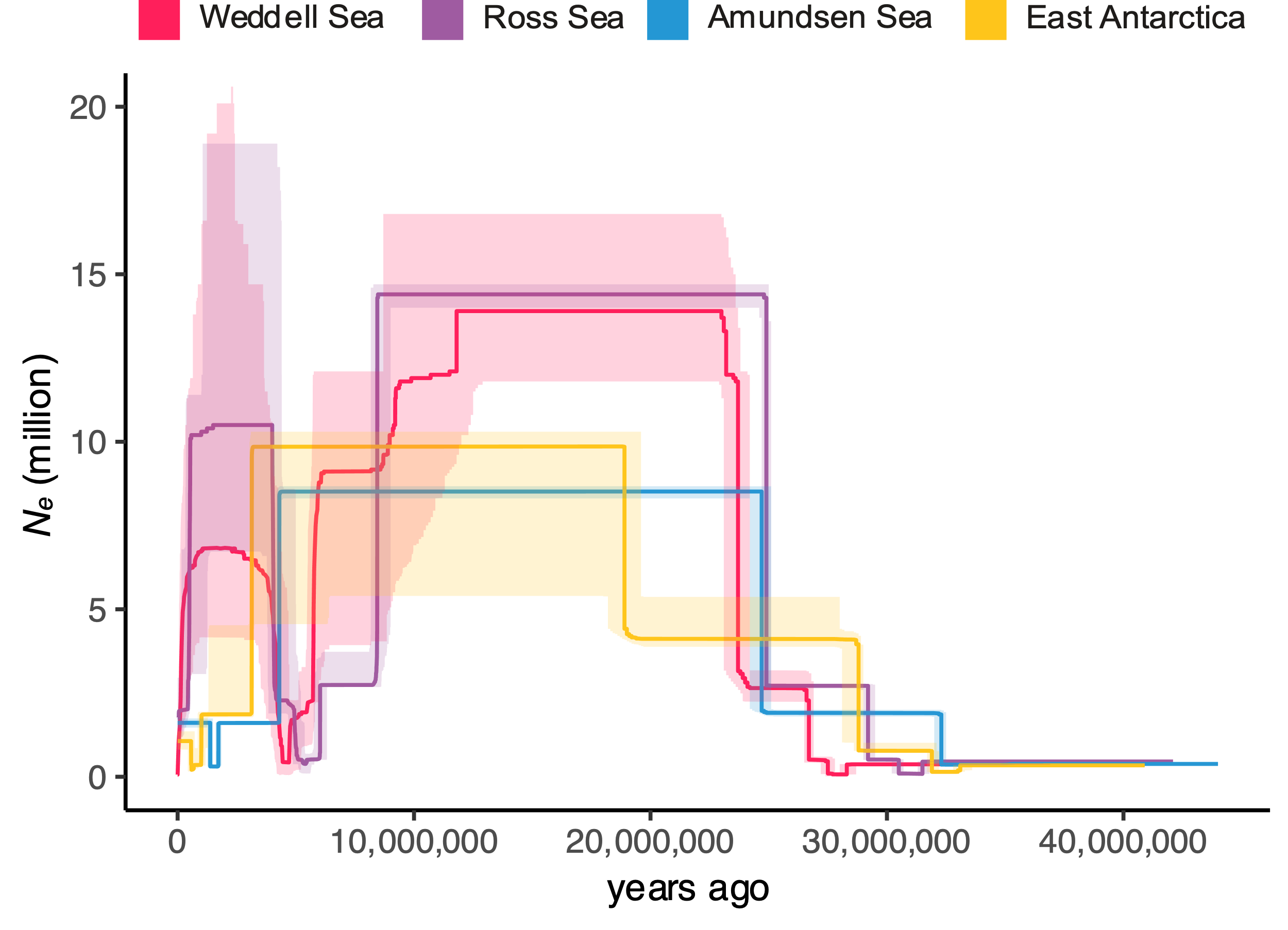

**Fig. S15.**

***StairwayPlot* reconstruction of past changes in effective population size over time in *Pareledone turqueti*.** Line = median, shaded area = 2.5% and 97.5% confidence limits. Data = 191,024-SNP dataset.

**Fig. S16.**

**Fit of the expected and observed one-dimensional (1D)-SFS under the best model evaluated (‘anc_psccc_fullcol2’) for *Pareledone turqueti*.** Marginal 1D-SFS of the observed data (black bars) is compared to the averaged expected SFS (light grey bars) obtained from 100 SFS approximated with 200,000 coalescent simulations. Error bars = range of the values obtained across 100 simulated expected SFS under the parameters that maximised the likelihoods. Data = 163,335-SNP dataset.

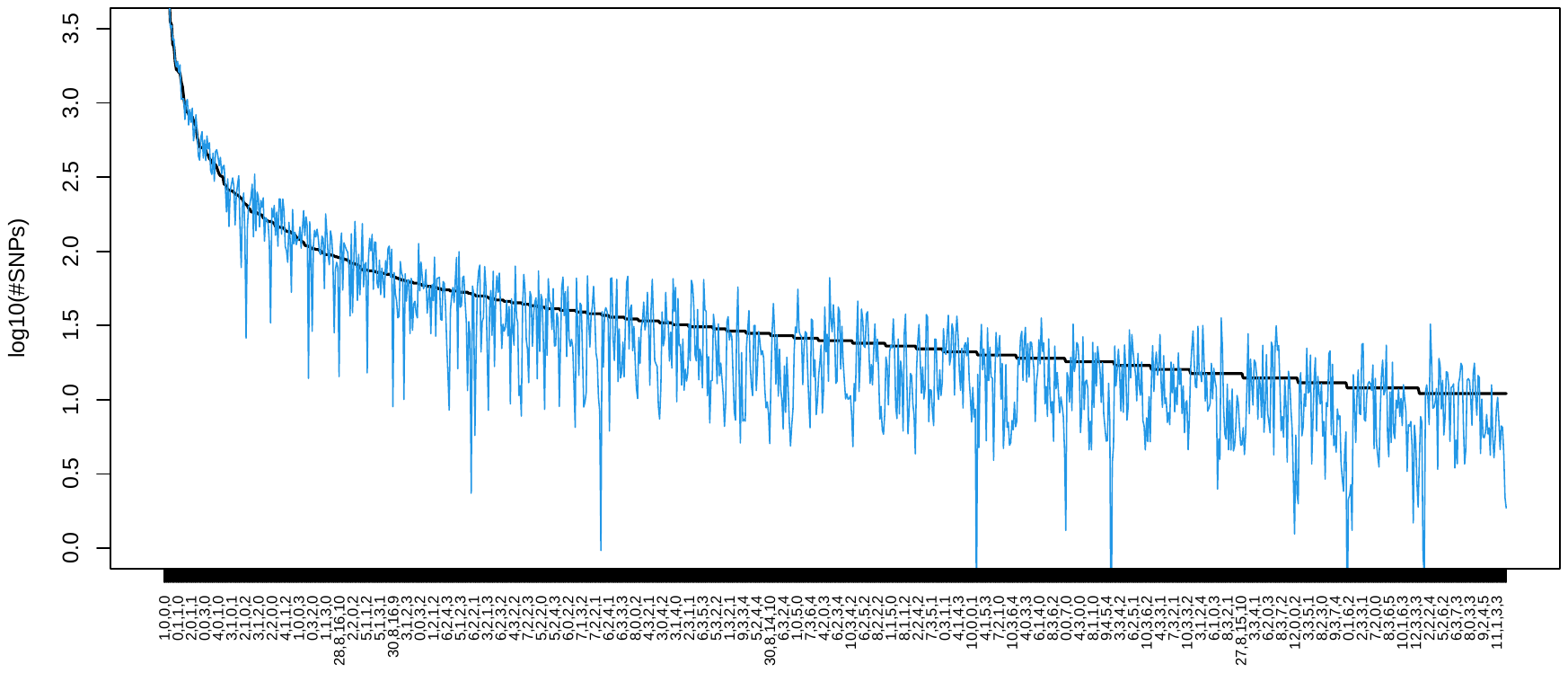

**Fig. S17.**

**Fit of the expected to observed four-dimensional (4D)-SFS under the best model evaluated (‘anc_psccc_fullcol2’) for *Pareledone turqueti*.** Only entries with more than 10 SNPs are shown. Entries in the x-axis are indicated by column in the format of (AS, RS, EA, WS), and numbers within each entry correspond to the count of the derived allele in Amundsen Sea (AS), Ross Sea (RS), East Antarctica (EA) and Weddell Sea (WS). Solid black line represents observed SFS, blue line represents averaged expected SFS. Averaged expected SFS was obtained from 100 SFS approximated with 200,000 coalescent simulations under the parameters that maximised the likelihoods. Data = 163,335-SNP dataset.

**Fig. S18.**

**Fit of the expected and observed one-dimensional (1D)-SFS under the step 2 model with no WAIS collapse (‘psccc_nocol’) for *Pareledone turqueti*.** Marginal 1D-SFS of the observed data (black bars) is compared to the averaged expected SFS (light grey bars) obtained from 100 SFS approximated with 200,000 coalescent simulations. Error bars = range of the values obtained across 100 simulated expected SFS under the parameters that maximised the likelihoods. Data = 163,335-SNP dataset.

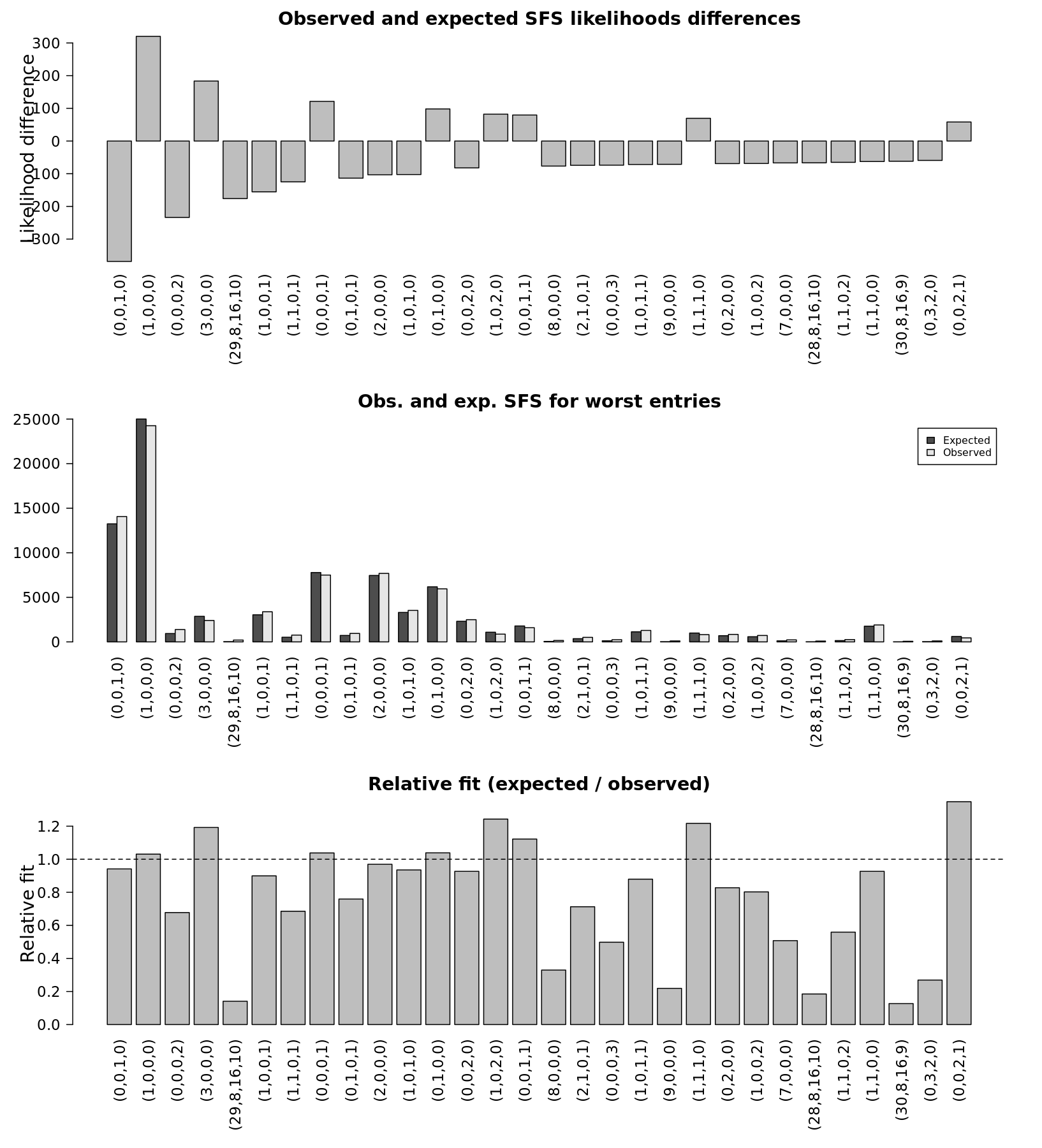

**Fig. S19.**

**Comparisons of the four-dimensional (4D)-joint observed and expected SFS for the 30 entries with the worst fit, under the step 2 model with no WAIS collapse (‘psccc_nocol’) for *Pareledone turqueti*.** Entries in the x-axis are indicated by column in the format of (WS, AS, RS, EA), and numbers within each entry correspond to the count of the derived allele in Weddell Sea (WS), Amundsen Sea (AS), Ross Sea (RS) and East Antarctica (EA). Averaged expected SFS was obtained from 100 SFS approximated with 200,000 coalescent simulations under the parameters that maximised the likelihoods. Top panel: comparison of the differences in likelihood (diff. lhood) between expected (exp) and observed (obs) SFS for all entries. Middle panel: comparison of the relative fit for all SFS entries. Bottom panel: Relative fit is defined as the relative number of SNP counts for a given entry (count of expected SNPs / count of observed SNPs). Data = 163,335-SNP dataset.

**Fig. S20.**

**Fit of the expected and observed one-dimensional (1D)-SFS under the step 2 model with partial WAIS collapse connecting Weddell Sea and Amundsen Sea (‘psccc_parcol’) for *Pareledone turqueti*.** Marginal 1D-SFS of the observed data (black bars) is compared to the averaged expected SFS (light grey bars) obtained from 100 SFS approximated with 200,000 coalescent simulations. Error bars = range of the values obtained across 100 simulated expected SFS under the parameters that maximised the likelihoods. Data = 163,335-SNP dataset.

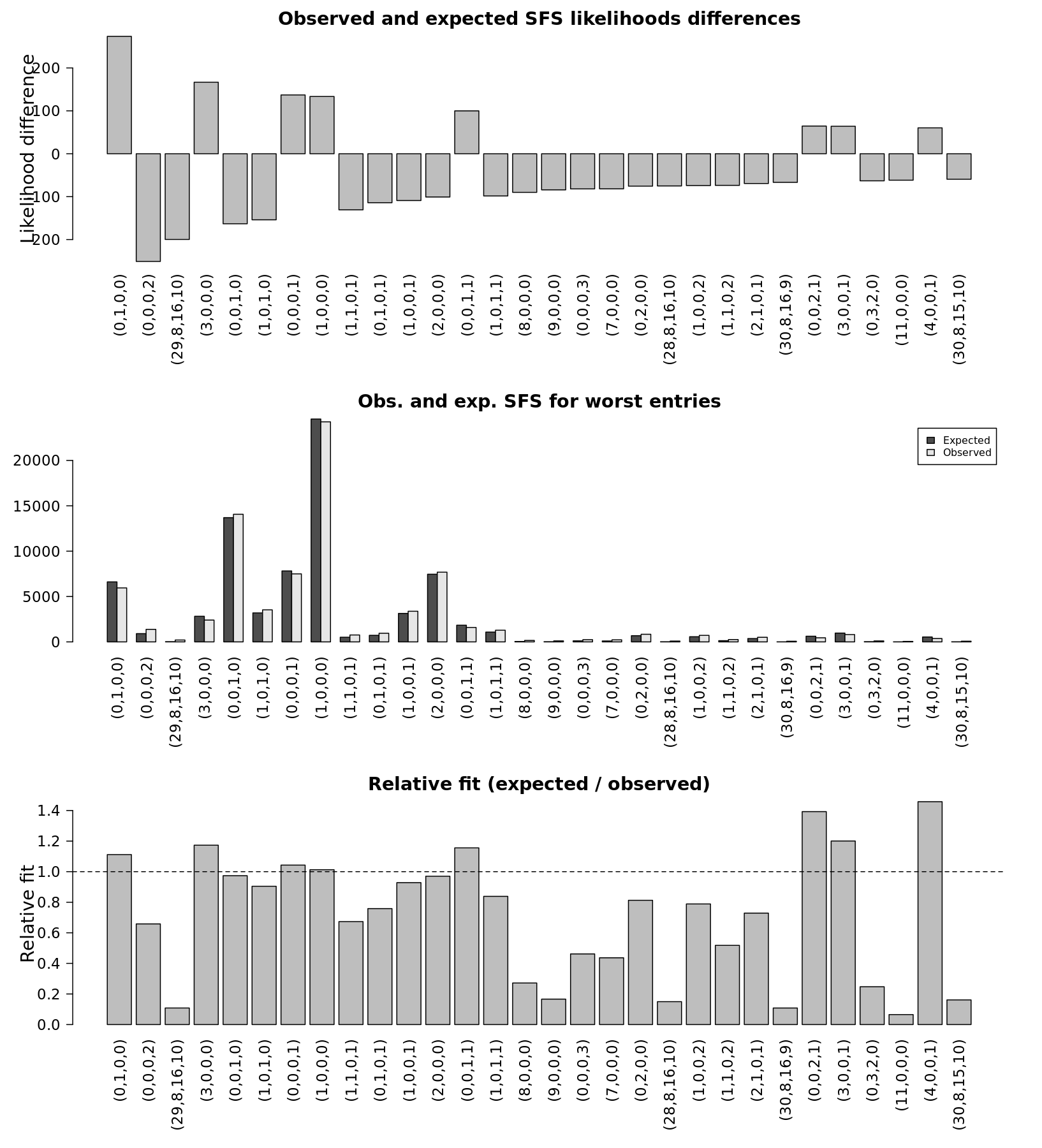

**Fig. S21.**

**Comparisons of the four-dimensional (4D)-joint observed and expected SFS for the 30 entries with the worst fit, under the step 2 model with partial WAIS collapse connecting Weddell Sea and Amundsen Sea (‘psccc_parcol’) for *Pareledone turqueti*.** Entries in the x-axis are indicated by column in the format of (WS, AS, RS, EA), and numbers within each entry correspond to the count of the derived allele in Weddell Sea (WS), Amundsen Sea (AS), Ross Sea (RS) and East Antarctica (EA). Averaged expected SFS was obtained from 100 SFS approximated with 200,000 coalescent simulations under the parameters that maximised the likelihoods. Top panel: comparison of the differences in likelihood (diff. lhood) between expected (exp) and observed (obs) SFS for all entries. Middle panel: comparison of the relative fit for all SFS entries. Bottom panel: Relative fit is defined as the relative number of SNP counts for a given entry (count of expected SNPs / count of observed SNPs). Data = 163,335-SNP dataset.

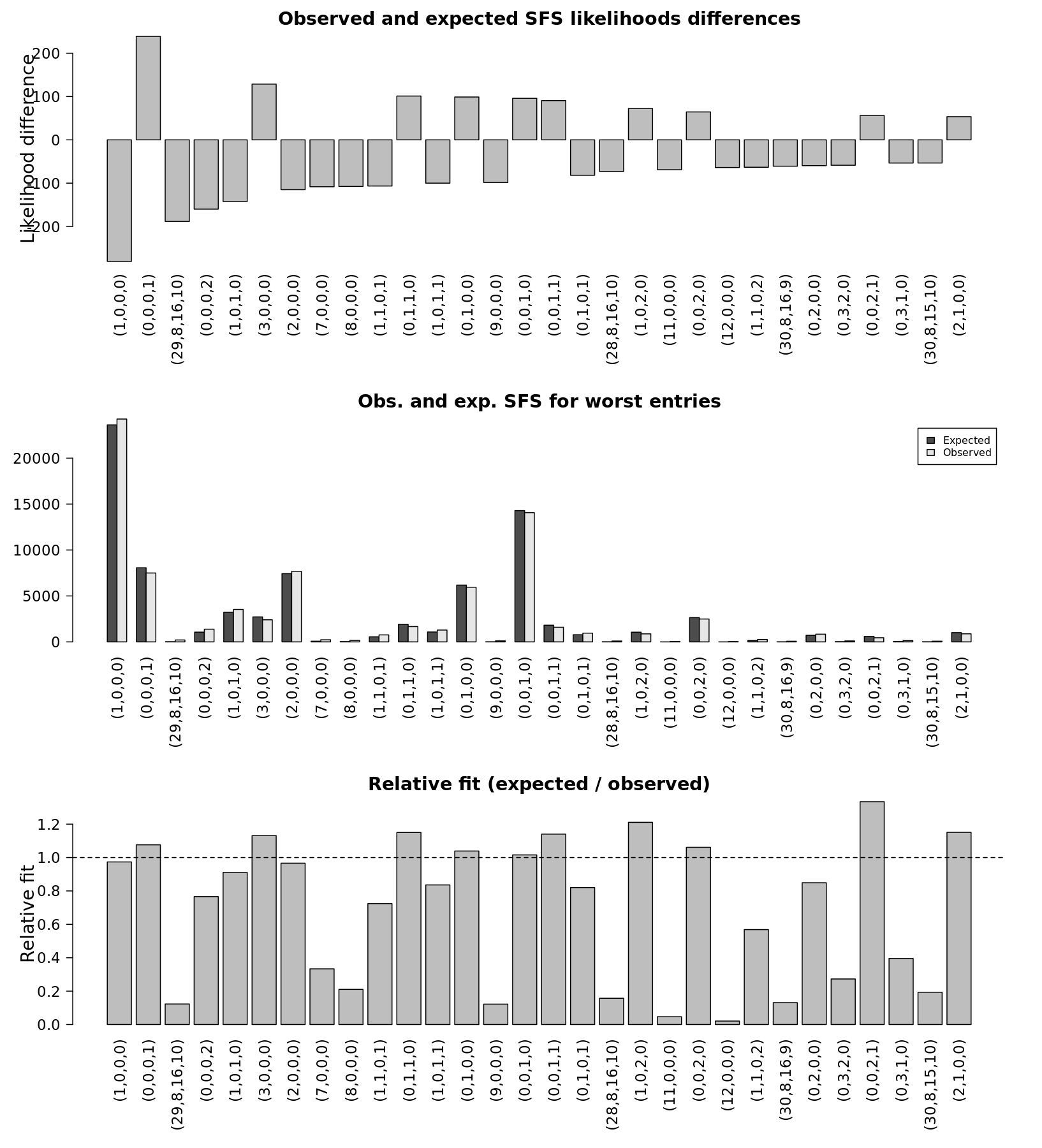
**Fig. S22.**

**Comparisons of the four-dimensional (4D)-joint observed and expected SFS for the 30 entries with the worst fit, under the step 3 complex model with complete WAIS collapse connecting Weddell Sea and Ross Sea, Weddell Sea and Amundsen Sea, and Amundsen Sea and Ross Sea (‘anc_psccc_fullcol2’) for *Pareledone turqueti*.** Entries in the x-axis are indicated by column in the format of (WS, AS, RS, EA), and numbers within each entry correspond to the count of the derived allele in Weddell Sea (WS), Amundsen Sea (AS), Ross Sea (RS) and East Antarctica (EA). Averaged expected SFS was obtained from 100 SFS approximated with 200,000 coalescent simulations under the parameters that maximised the likelihoods. Top panel: comparison of the differences in likelihood (diff. lhood) between expected (exp) and observed (obs) SFS for all entries. Middle panel: comparison of the relative fit for all SFS entries. Bottom panel: Relative fit is defined as the relative number of SNP counts for a given entry (count of expected SNPs / count of observed SNPs). Data = 163,335-SNP dataset.

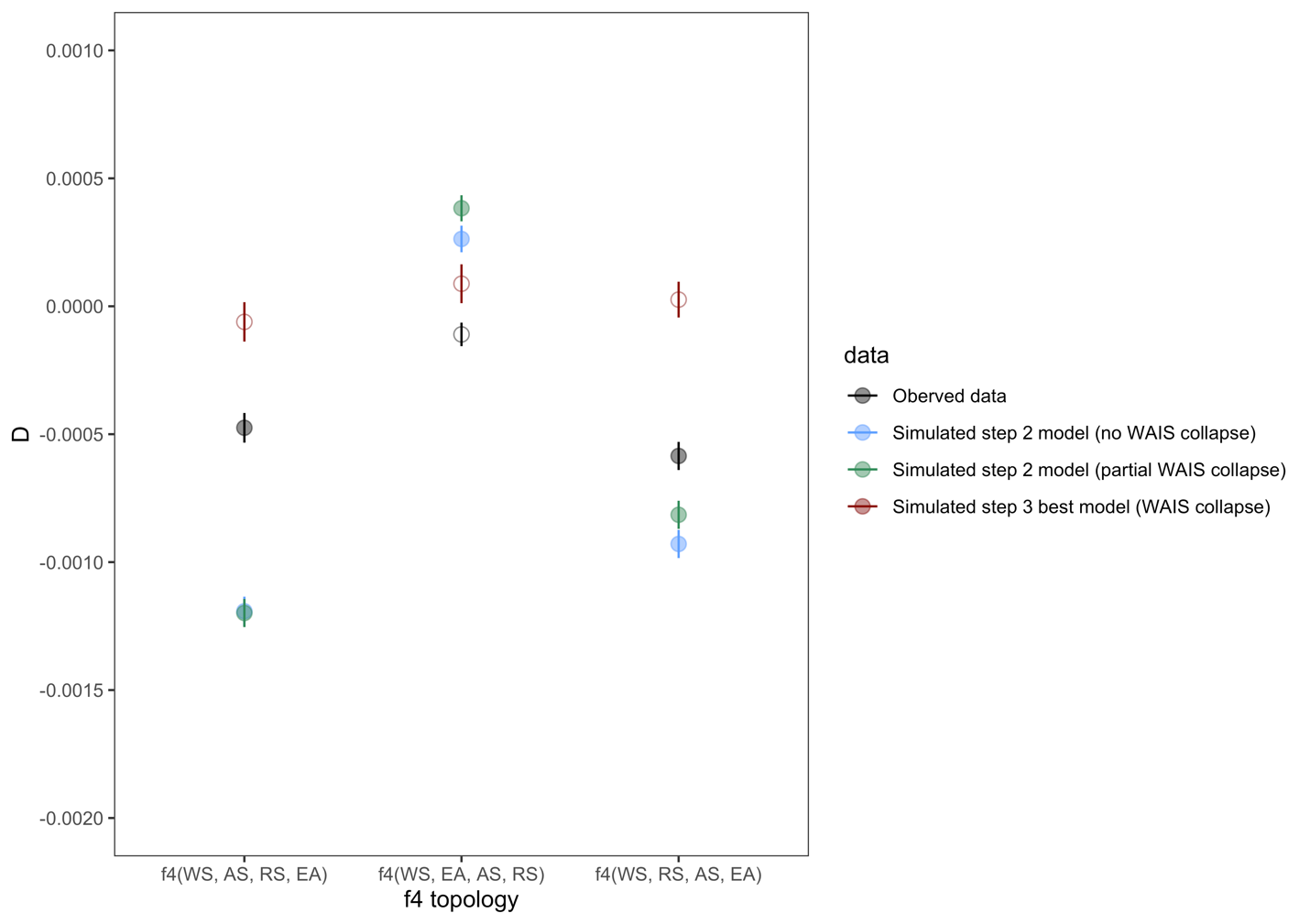

**Fig. S23.**

**Results of *f*4 statistic using 100,000 SNPs simulated under the best fit *fastsimcoal* model with full WAIS collapse (“anc_psccc_fullcol1”) and the worst fit simpler step 2 model with partial (“psccc_parcol”) or no (“psccc_nocol”) WAIS collapse.** Simulated data was generated with maximum likelihood parameters. Observed data was calculated from the observed SNPs used in *fastsimcoal* (Data = 163,335-SNP dataset). Abbreviations: WS (Weddell Sea), AS (Amundsen Sea), RS (Ross Sea) and EA (East Antarctica). Error bars (vertical lines) = standard errors, filled circles = significant (Z-score values > 3 or < -3).

**Fig. S24.**

**Redundancy analysis (RDA) showing genotype–environment association in *Pareledone turqueti* samples across the Scotia Sea (n = 52) on the first two constrained axes.** Grey dots represent SNPs, and coloured circles represent individual sample defined by geographical locations. Vectors represent environmental predictors, including water depth, salinity, nitrate, temperature and longitude. Data = 5,188-SNP dataset.

**Fig. S25.**
**Clustering analysis using *Structure* inferred *K* = 7 for *P. turqueti*.** Each horizontal bar represents an individual sample, bars are grouped by geographical locations, colours within each bar correspond to the proportion of each genetic cluster in the individual. Individual islands within the South Shetland Islands group are labelled separately. Data = 5,188-SNP dataset.

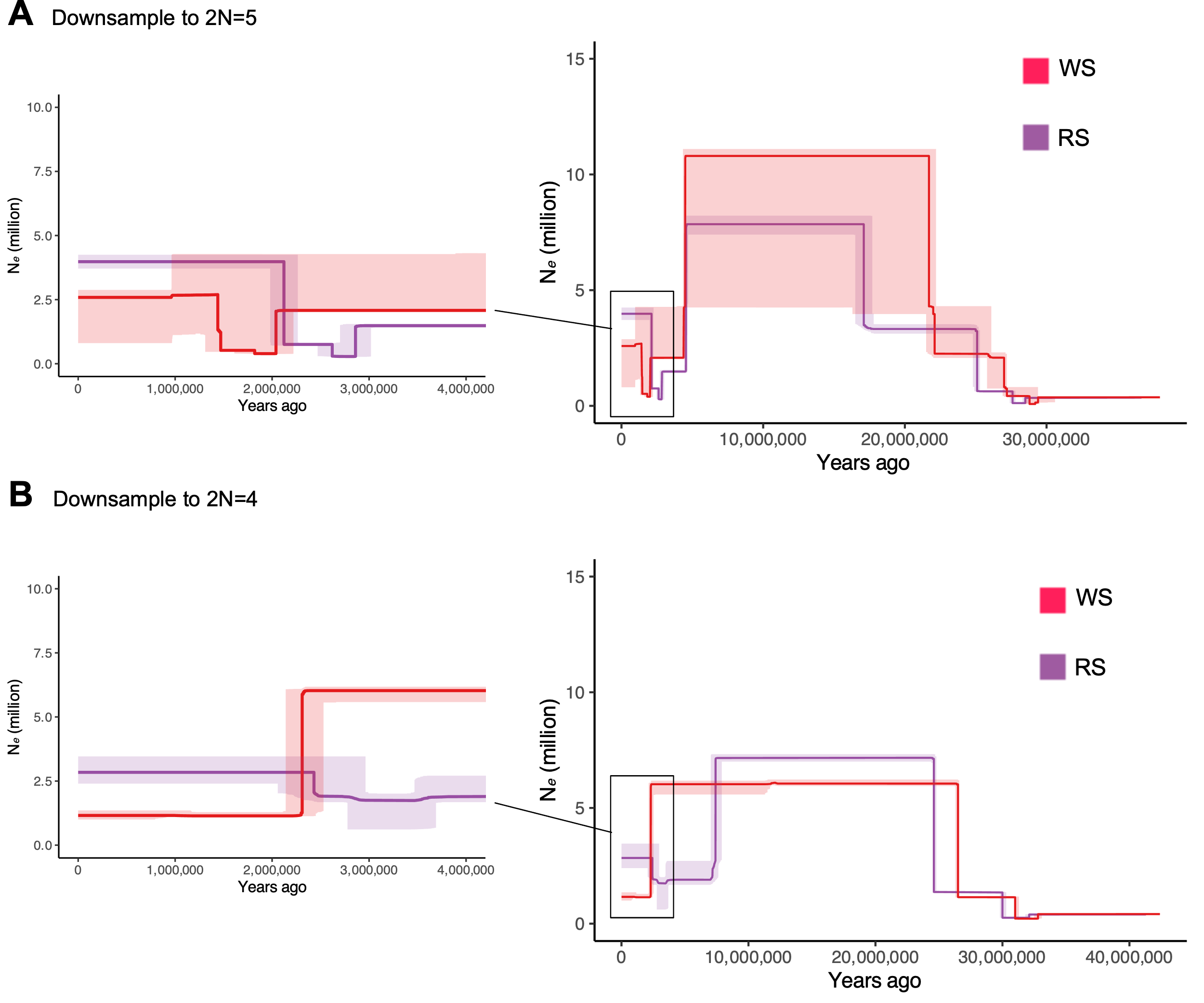

**Fig. S26.**

***StairwayPlot* reconstruction of past changes in effective population size over time in *P. turqueti*.** Weddell Sea (WS) and Ross Sea (RS) samples were downsampled to 2N = 5 (**A**) and 2N = 4 (**B**) to examine the influence of small sample size in *StairwayPlot*. Data = 191,024-SNP dataset.

**Table S1.
Genetic diversity statistics (expected and observed heterozygosity (H_e_, H_o_) and Inbreeding coefficient (F_IS_) across individuals per sampled location in *Pareledone turqueti*.** Data = 5,188-SNP dataset.

| **Location** | **Observed heterozygosity (H_o_)** | **Expected Heterozygosity (H_e_)** | **Inbreeding coefficient (F_IS_)** |
| --- | --- | --- | --- |
| Shag Rocks | 0.248 | 0.204 | -0.218 |
| South Georgia | 0.260 | 0.213 | -0.223 |
| Elephant Is. | 0.213 | 0.191 | -0.114 |
| South Shetland Is. | 0.214 | 0.196 | -0.094 |
| Amundsen Sea | 0.217 | 0.189 | -0.145 |
| South Weddell Sea | 0.202 | 0.193 | -0.048 |
| East Weddell Sea | 0.213 | 0.187 | -0.141 |
| Ross Sea | 0.203 | 0.180 | -0.128 |
| Adélie Land | 0.211 | 0.182 | -0.161 |
| Prydz Bay | 0.229 | 0.202 | -0.133 |

Table S2.
Demographic parameters inferred in the step 3 best model (anc_psccc_fullcol2) in *Pareledone turqueti*. Maximum-likelihood (ML) parameter estimates were extracted from the best run with the highest composite likelihood among 100 replicates. Migration from pop i to pop j is denoted as MIGij looking backward in time. Number of migrants per generation from pop j to pop i is denoted as IM_MIGij$, and is scaled as 2Nm (2N = population effective sizes in diploid), looking forward in time. Effective population sizes are given in the number of haploids (N). Estimations of timing of events are given in the number of generations (gen) and years. The 95% confidence intervals (CI) were generated from 100 nonparametric bootstrapped datasets. Data = 163,335-SNP dataset. Abbreviations: Weddell Sea (WS), Amundsen Sea (AS), Ross Sea (RS), East Antarctica (EA).

|  |  |  | **95% CI** | |
| --- | --- | --- | --- | --- |
| **Parameter** | **Parameter description** | **ML estimate** | **Lower bound** | **Upper bound** |
| NEA$ | Effective population size of EA at T3 | 88644 | 36132 | 106422 |
| NWS$ | Effective population size of WS at T3 | 935567 | 901972 | 1401818 |
| NAS$ | Effective population size of AS at T3 | 57021 | 25702 | 72557 |
| NRS$ | Effective population size of RS at T3 | 1060714 | 1053829 | 1454709 |
| NEAC$ | Effective population size of EA at T2 | 498346 | 537748 | 641332 |
| NWSC$ | Effective population size of WS at T2 | 3060169 | 3011287 | 3306845 |
| NASC$ | Effective population size of AS at T2 | 632696 | 734274 | 862138 |
| NRSC$ | Effective population size of RS at T2 | 5073835 | 3818690 | 4314333 |
| NANC3$ | Effective population size of the ancestral population of AS, WS, RS and EA between T1 and T2 | 862974 | 799336 | 880704 |
| NANC2$ | Effective population size of the ancestral population of AS, WS, RS and EA between T0 and T1 | 8670493 | 8777970 | 9013260 |
| NANC1$ | Effective population size of the ancestral population of AS, WS, RS and EA before T0 | 401522 | 350740 | 362583 |
| T0 (gen) | Time of first demographic change in the ancestral population of WS, AS, RS and EA (in generations) | 2764388 | 2763732 | 2784210 |
| T1 (gen) | Time of second demographic change in the ancestral population of WS, AS, RS and EA (in generations) | 253663 | 293378 | 304923 |
| T2 (gen) | Time of trans-west Antarctic seaway connectivity between WS-AS-RS begins (in generations) | 250989 | 288208 | 299592 |
| T3 (gen) | Time of trans-west Antarctic seaway connectivity between WS-AS-RS ceases; time of contemporary gene flow between WS-AS-RS-EA along circumpolar current begins (in generations) | 7232 | 4480 | 11600 |
| T0 (year) | Time of first demographic change in the ancestral population of WS, AS, RS and EA (in years) | 33172656 | 33164784 | 33410520 |
| T1 (year) | Time of second demographic change in the ancestral population of WS, AS, RS and EA (in years) | 3043956 | 3520536 | 3659076 |
| T2 (year) | Time of trans-west Antarctic seaway connectivity between WS-AS-RS begins (in years) | 3011868 | 3458496 | 3595104 |
| T3 (year) | Time of trans-west Antarctic seaway connectivity between WS-AS-RS ceases; time of contemporary gene flow between WS-AS-RS-EA along circumpolar current begins (in years) | 86784 | 53760 | 139200 |
| MIG10$ | Migration rate from AS to WS at T3 | 1.13E-04 | 5.75E-05 | 1.51E-04 |
| MIG30$ | Migration rate from EA to WS at T3 | 1.91E-04 | 1.48E-04 | 2.91E-04 |
| MIG01$ | Migration rate from WS to AS at T3 | 5.38E-07 | 0.00E+00 | 9.99E-07 |
| MIG21$ | Migration rate from RS to AS at T3 | 6.15E-10 | 0.00E+00 | 3.71E-08 |
| MIG12$ | Migration rate from AS to RS at T3 | 3.27E-04 | 2.45E-04 | 5.24E-04 |
| MIG32$ | Migration rate from EA to RS at T3 | 1.66E-04 | 1.36E-04 | 2.77E-04 |
| MIG03$ | Migration rate from WS to EA at T3 | 1.34E-05 | 4.24E-06 | 6.74E-06 |
| MIG23$ | Migration rate from RS to EA at T3 | 8.50E-07 | 0.00E+00 | 1.09E-06 |
| MIG10C$ | Migration rate from AS to WS at T2 | 1.84E-07 | 0.00E+00 | 1.81E-08 |
| MIG20C$ | Migration rate from RS to WS at T2 | 2.06E-08 | 0.00E+00 | 9.91E-08 |
| MIG01C$ | Migration rate from WS to AS at T2 | 3.05E-06 | 2.20E-06 | 2.60E-06 |
| MIG21C$ | Migration rate from RS to AS at T2 | 3.39E-06 | 3.29E-06 | 3.78E-06 |
| MIG02C$ | Migration rate from WS to RS at T2 | 4.65E-06 | 1.86E-06 | 3.44E-06 |
| MIG12C$ | Migration rate from AS to RS at T2 | 6.21E-07 | 8.57E-09 | 9.52E-08 |
| IM_MIG10$ | Number of migrants from WS to AS at T3 | 3.23 | 0.74 | 5.48 |
| IM_MIG30$ | Number of migrants from WS to EA at T3 | 8.47 | 2.67 | 15.47 |
| IM_MIG01$ | Number of migrants from AS to WS at T3 | 0.25 | 0E+00 | 0.70 |
| IM_MIG21$ | Number of migrants from AS to RS at T3 | 3.26E-04 | 0E+00 | 0.03 |
| IM_MIG12$ | Number of migrants from RS to AS at T3 | 9.34 | 3.15 | 19.00 |
| IM_MIG32$ | Number of migrants from RS to EA at T3 | 7.35 | 2.46 | 14.76 |
| IM_MIG03$ | Number of migrants from EA to WS at T3 | 6.27 | 1.91 | 4.72 |
| IM_MIG23$ | Number of migrants from EA to RS at T3 | 0.45 | 0E+00 | 0.79 |
| IM_MIG10C$ | Number of migrants from WS to AS at T2 | 0.06 | 0E+00 | 0.01 |
| IM_MIG20C$ | Number of migrants from WS to RS at T2 | 0.05 | 0E+00 | 0.21 |
| IM_MIG01C$ | Number of migrants from AS to WS at T2 | 4.67 | 3.31 | 4.30 |
| IM_MIG21C$ | Number of migrants from AS to RS at T2 | 8.60 | 6.28 | 8.15 |
| IM_MIG02C$ | Number of migrants from RS to WS at T2 | 7.12 | 2.81 | 5.68 |
| IM_MIG12C$ | Number of migrants from RS to AS at T2 | 0.20 | 3E-03 | 0.04 |

**Table S3.**

**Comparisons of simple demographic models with three populations, with alternative tree topologies.** (**A**) Models tested between Weddell Sea (WS), Ross Sea (RS), East Antarctica (EA). Data = 167,612-SNP dataset. (**B**) Models tested between WS, RS and South Shetland Islands (SHE). Data = 195,003-SNP dataset. Model label corresponds to model label in Fig. S13. Delta AIC and relative likelihoods were calculated following Excoffier et al. (2013) (*36*). Abbreviations: Lhood = log likelihoods, AIC = Akaike Information Criterion.

| **(A)** | Model name | Lhood | Number of parameters | AIC | Delta AIC | Relative likelihood (Akaike's weight of evidence) |
| --- | --- | --- | --- | --- | --- | --- |
|  | WS_RS_EA_fullcol1 | -532682.74 | 11 | 2453558.72 | 0.00 | 1 |
|  | WS_RS_EA | -532730.82 | 5 | 2453768.18 | 209.46 | 0 |
|  | EA_WSRS | -538467.75 | 6 | 2480194.47 | 26635.75 | 0 |
| **(B)** |  |  |  |  |  |  |
|  | WS_RS_SHE | -627444.94 | 5 | 2890021.40 | 0.00 | 1 |
|  | WS_RS_SHE_fullcol1 | -627445.57 | 11 | 2890036.31 | 14.91 | 0 |
|  | SHE_WSRS | -627557.11 | 6 | 2890540.03 | 518.63 | 0 |

**Table S4**.

**Comparisons of simple demographic models with three populations with alternative tree topologies, with present day gene flow linked to the Antarctic circumpolar current (clockwise) and Antarctic slope current (counter-clockwise).** (**A**) Models tested between Weddell Sea (WS), Ross Sea (RS), East Antarctica (EA). Data = 167,612-SNP dataset. (**B**) Models tested between WS, RS and South Shetland Islands (SHE). Data = 195,003-SNP dataset. Model label corresponds to model label in Fig. S14.. Delta AIC and relative likelihoods were calculated following Excoffier et al. (2013) (*36*). Abbreviations: Lhood = log likelihoods, AIC = Akaike Information Criterion.

| **(A)** | Model name | Lhood | Number of parameters | AIC | Delta AIC | Relative likelihood (Akaike's weight of evidence) |
| --- | --- | --- | --- | --- | --- | --- |
|  | WS_RS_EA_fullcol1_cc | -530471.00 | 15 | 2443379.42 | 0.00 | 1 |
|  | EA_WSRS_cc | -530731.89 | 10 | 2444571.10 | 1191.68 | 0 |
| **(B)** |  |  |  |  |  |  |
|  | WS_RS_SHE_fullcol1_cc | -624560.49 | 15 | 2876755.63 | 0.00 | 1 |
|  | SHE_WSRS_cc | -624903.07 | 10 | 2878323.55 | 1567.92 | 0 |

Table S5.

**Results of outgroup-ƒ3-statistics between pairs of populations.** As ƒ3 value increases, more derived allele frequency is shared between population A and population B related to an outgroup population (population C). Abbreviations: Weddell Sea (WS), South Shetland Islands (SHE), Amundsen Sea (AS), Ross Sea (RS), East Antarctica (EA), Shag Rocks and South Georgia (SGSR; samples combined). Z-score values > 3 or < -3 = significance, stderr = standard error, nsnps = number of SNPs involved in the statistics. Data = 120,857-SNP dataset.

| **population A** | **population B** | **population C** | **ƒ3** | **stderr** | **Zscore** | **nsnps** |
| --- | --- | --- | --- | --- | --- | --- |
| AS | RS | SRSG | 0.054015 | 0.001107 | 48.80 | 92000 |
| RS | EA | SRSG | 0.046025 | 0.001025 | 44.92 | 93586 |
| RS | WS | SRSG | 0.045840 | 0.000870 | 52.71 | 105128 |
| RS | SHE | SRSG | 0.043447 | 0.000787 | 55.24 | 97556 |
| AS | EA | SRSG | 0.043384 | 0.000957 | 45.35 | 86461 |
| AS | SHE | SRSG | 0.042471 | 0.000841 | 50.47 | 90359 |
| AS | WS | SRSG | 0.042254 | 0.000847 | 49.88 | 99122 |
| WS | SHE | SRSG | 0.039757 | 0.000702 | 56.63 | 102545 |
| WS | EA | SRSG | 0.039279 | 0.000876 | 44.84 | 99159 |
| SHE | EA | SRSG | 0.030848 | 0.000714 | 43.21 | 92090 |

**Table S6.**

**Results of *D*-statistic between (in the form of BABA-ABBA) examining patterns of alleles sharing across four populations, and indicates whether there is excess allele sharing between distinct populations.** Top panel: *D*-statistic is presented in the form of *D*(Pop, SHE, WS, Out), which examines whether there is excess allele sharing between SHE and WS (D < 0; ABBA) or Pop and WS (*D* > 0; BABA). Bottom panel: *D*-statistic is presented in the form of *D*(Pop, EA, WS, Out), which examines whether there is excess allele sharing between EA and WS (*D* < 0; ABBA) or Pop and WS (*D* > 0; BABA). Abbreviations: Weddell Sea (WS), South Shetland Islands (SHE), Amundsen Sea (AS), Ross Sea (RS), East Antarctica (EA), Shag Rocks and South Georgia (SRSG; samples combined as an outgroup population [Out]). Z-score values > 3 or < -3 = significance, stderr = standard error, nsnps = number of SNPs involved in the statistics. Data = 120,857-SNP dataset.

| ***D*-statistic presentation** | **W** | **X** | **Y** | **Z** | ***D*** | **stderr** | **Zscore** | **BABA** | **ABBA** | **nsnps** |
| --- | --- | --- | --- | --- | --- | --- | --- | --- | --- | --- |
| *D*(Pop, SHE, WS, Out)  / *D*(W, X, Y, Z) | WS | SHE | WS | SRSG | 0.0113 | 0.001857 | 6.084 | 1576 | 1541 | 120857 |
|  | RS | SHE | WS | SRSG | 0.0284 | 0.001838 | 15.447 | 1554 | 1468 | 120857 |
| *D*(Pop, EA, WS, Out)  / *D*(W, X, Y, Z) | WS | EA | WS | SRSG | 0.0130 | 0.002007 | 6.465 | 1638 | 1596 | 120857 |
|  | RS | EA | WS | SRSG | 0.0295 | 0.001870 | 15.77 | 1616 | 1523 | 120857 |

**Table S7.**

**Summary of likelihoods for the model tested at step 1 in *Pareledone turqueti*.** Model label corresponds to model label in fig. S3-S4. Delta AIC and relative likelihoods were calculated following Excoffier et al. (2013) (*36*). Abbreviations: Lhood = log likelihoods, AIC = Akaike Information Criterion. Data = 163,335-SNP dataset.

| **Model label** | **Lhood** | **Number of parameters** | **AIC** | **Delta AIC** | **Relative likelihood (Akaike's weight of evidence)** |
| --- | --- | --- | --- | --- | --- |
| psc_nocol | -548951.53 | 17 | 2528504.77 | 0.00 | 1.00 |
| psc_parcol | -548960.42 | 19 | 2528549.68 | 44.91 | 0.00 |
| psc_fullcol2 | -548969.37 | 23 | 2528598.91 | 94.14 | 0.00 |
| psc_parcol2 | -548987.64 | 19 | 2528675.09 | 170.32 | 0.00 |
| psc_fullcol1 | -548988.68 | 19 | 2528679.88 | 175.11 | 0.00 |
| psc_conflow | -549019.38 | 12 | 2528807.25 | 302.48 | 0.00 |

**Table S8.**

**Summary of likelihoods for the model tested at Step 2 in *Pareledone turqueti*.** Model label corresponds to model label in Fig. S3-S4. Delta AIC and relative likelihoods were calculated following Excoffier et al. (2013) (*36*). Abbreviations: Lhood = log likelihoods, AIC = Akaike Information Criterion. Data = 163,335-SNP dataset.

| **Model label** | **Lhood** | **Number of parameters** | **AIC** | **Delta AIC** | **Relative likelihood (Akaike's weight of evidence)** |
| --- | --- | --- | --- | --- | --- |
| psccc_fullcol2 | -548629.82 | 27 | 2527042.97 | 0.00 | 1.00 |
| psccc_nocol | -548635.01 | 21 | 2527054.85 | 11.88 | 0.00 |
| psccc_parcol | -548640.40 | 23 | 2527083.70 | 40.73 | 0.00 |
| psccc_fullcol3 | -548651.19 | 25 | 2527137.38 | 94.41 | 0.00 |
| psccc_fullcol1 | -548652.25 | 23 | 2527138.28 | 95.31 | 0.00 |
| psccc_fullcol4 | -548652.07 | 25 | 2527141.45 | 98.48 | 0.00 |
| psccc_parcol2 | -548668.16 | 23 | 2527211.55 | 168.58 | 0.00 |
| psccc_conflow | -548754.62 | 16 | 2527595.80 | 552.83 | 0.00 |

**Table S9.**

**Summary of likelihoods for the model tested at Step 3 in *Pareledone turqueti*.** Model label corresponds to model label in Fig. S3. Delta AIC and relative likelihoods were calculated following Excoffier et al. (2013) (*36*). Abbreviations: Lhood = log likelihoods, AIC = Akaike Information Criterion. Data = 163,335-SNP dataset.

| **Model label** | **Lhood** | **Number of parameters** | **AIC** | **Delta AIC** | **Relative likelihood (Akaike's weight of evidence)** |
| --- | --- | --- | --- | --- | --- |
| anc_psccc_fullcol2 | -548592.40 | 29 | 2526874.60 | 0.00 | 1.00 |
| anc_psccc_parcol | -548638.55 | 25 | 2527079.16 | 204.57 | 0.00 |
| anc_psccc_nocol | -548640.51 | 23 | 2527084.18 | 209.58 | 0.00 |

**Table S10.**
**Alternative estimations of demographic event timing in the best model (anc_psccc_fullcol2) in *Pareledone turqueti*.** Lower estimate of 11 years was inferred from *Pareledone charcoti*’s age estimation), upper estimate of 13 years was inferred from the upper limit of *P. turqueti* age estimation.

|  |  | Generation time assumption (years) | | |
| --- | --- | --- | --- | --- |
| Parameter label | Parameter description | 12 (used in manuscript) | 11 (lower estimate) | 13 (upper estimate) |
| T0 | Time of first demographic change in the ancestral population of WS, AS, RS and EA (in years) | 33172656 | 30408268 | 35937044 |
| T1 | Time of second demographic change in the ancestral population of WS, AS, RS and EA (in years) | 3043956 | 2790293 | 3297619 |
| T2 | Time of trans-west Antarctic seaway connectivity between WS-AS-RS begins (in years) | 3011868 | 2760879 | 3262857 |
| T3 | Time of trans-west Antarctic seaway connectivity between WS-AS-RS ceases; time of contemporary gene flow between WS-AS-RS-EA linked to circumpolar current begins (in years) | 86784 | 79552 | 94016 |

**Table S11.**

**Demographic parameters inferred in the best simple three population model between WS, RS and EA (WS_RS_EA_fullcol1_cc) in *Pareledone turqueti*.** Maximum-likelihood (ML) parameter estimates were extracted from the best run with the highest composite likelihood among 100 replicates. Migration rate (m) from pop i to pop j is denoted as MIGij looking backward in time. Number of migrants per generation from pop j to pop i is denoted as IM_MIGij$, and is scaled as 2Nm (2N = population effective sizes in diploid), looking forward in time. Effective population sizes are given in the number of haploids (N). Estimations of timing of events are given in the number of generations (gen) and years (assuming a generation time of 12 years). Data = 167,612-SNP dataset. Abbreviations: Weddell Sea (WS), Ross Sea (RS), East Antarctica (EA).

| **Parameter** | **ML estimate** | **Parameter description** |
| --- | --- | --- |
| NEA$ | 183675 | Effective population size of EA at T2 |
| NWS$ | 1112944 | Effective population size of WS at T2 |
| NRS$ | 776081 | Effective population size of RS at T2 |
| NEAC$ | 3059380 | Effective population size of EA at T1 |
| NWSC$ | 7071085 | Effective population size of WS at T1 |
| NRSC$ | 6811239 | Effective population size of RS at T1 |
| NANC$ | 3100488 | Effective population size of the ancestral population of WS, RS and EA before T1 |
| T1 (gen) | 329657 | Time of T1 event (in generation) |
| T2 (gen) | 9632 | Time of T2 event (in generation) |
| T1 (year) | 3955884 | Time of T1 event (in year) |
| T2 (year) | 115584 | Time of T2 event (in year) |
| MIG10C$ | 1.18E-06 | Migration rate from RS to WS at T1 |
| MIG01C$ | 2.25E-06 | Migration rate from WS to RS at T1 |
| MIG20$ | 1.02E-04 | Migration rate from EA to WS at T2 |
| MIG21$ | 7.88E-05 | Migration rate from EA to RS at T2 |
| MIG02$ | 1.74E-05 | Migration rate from WS to EA at T2 |
| MIG12$ | 2.05E-05 | Migration rate from RS to EA at T2 |
| IM_MIG10C$ | 4.03 | Number of migrants per generation from WS to RS at T1 |
| IM_MIG01C$ | 7.95 | Number of migrants per generation from RS to WS at T1 |
| IM_MIG20$ | 0.21 | Number of migrants per generation from WS to EA at T2 |
| IM_MIG21$ | 7.24 | Number of migrants per generation from RS to EA at T2 |
| IM_MIG02$ | 9.67 | Number of migrants per generation from EA to WS at T2 |
| IM_MIG12$ | 7.94 | Number of migrants per generation from EA to RS at T2 |

**Table S12.**

**Demographic parameters inferred in the best simple three population model between WS, RS and SHE (WS_RS_SHE_fullcol1) in *Pareledone turqueti*.** Maximum-likelihood (ML) parameter estimates were extracted from the best run with the highest composite likelihood among 100 replicates. Migration rate (m) from pop i to pop j is denoted as MIGij looking backward in time. Number of migrants per generation from pop j to pop i is denoted as IM_MIGij$, and is scaled as 2Nm (2N = population effective sizes in diploid), looking forward in time. Effective population sizes are given in the number of haploids (N). Estimations of timing of events are given in the number of generations (gen) and years (assuming a generation time of 12 years). Data = 195,003-SNP dataset. Abbreviations: Weddell Sea (WS), Ross Sea (RS), South Shetland Islands (SHE).

| **Parameter** | **ML estimate** | **Parameter description** |
| --- | --- | --- |
| NSHE$ | 716329 | Effective population size of SHE at T2 |
| NWS$ | 2573301 | Effective population size of WS at T2 |
| NRS$ | 1463931 | Effective population size of RS at T2 |
| NSHEC$ | 4106990 | Effective population size of SHE at T1 |
| NWSC$ | 7059195 | Effective population size of WS at T1 |
| NRSC$ | 6373303 | Effective population size of RS at T1 |
| NANC$ | 3225487 | Effective population size of the ancestral population of WS, RS and SHE before T1 |
| T1 (gen) | 329512 | Time of T1 event (in generation) |
| T2 (gen) | 23502 | Time of T2 event (in generation) |
| T1 (year) | 3954144 | Time of T1 event (in year) |
| T2 (year) | 282024 | Time of T2 event (in year) |
| MIG10C$ | 1.53E-06 | Migration rate from RS to WS at T1 |
| MIG01C$ | 2.49E-06 | Migration rate from WS to RS at T1 |
| MIG20$ | 2.61E-05 | Migration rate from SHE to WS at T2 |
| MIG21$ | 7.87E-06 | Migration rate from SHE to RS at T2 |
| MIG02$ | 7.17E-06 | Migration rate from WS to SHE at T2 |
| MIG12$ | 1.06E-05 | Migration rate from RS to SHE at T2 |
| IM_MIG10C$ | 4.88 | Number of migrants per generation from WS to RS at T1 |
| IM_MIG01C$ | 8.78 | Number of migrants per generation from RS to WS at T1 |
| IM_MIG20$ | 9.37 | Number of migrants per generation from WS to SHE at T2 |
| IM_MIG21$ | 2.82 | Number of migrants per generation from RS to SHE at T2 |
| IM_MIG02$ | 9.23 | Number of migrants per generation from SHE to WS at T2 |
| IM_MIG12$ | 7.77 | Number of migrants per generation from SHE to RS at T2 |

**Table S13.
Results of Generalised Dissimilarity Modelling (GDM) including model fit and relative importance of geographic distance and sample collection depth on explaining the genetic differentiation across *P. turqueti* samples. (A)** at a circumpolar level (n = 93), and **(B)** within the Scotia Sea (n = 52). Importance was defined by the % in deviance explained by the full model after permutation. Significance was determined with 999 permutations with significance level at 0.05. Data = 120,857-SNP dataset.

| **(A)** *Pareledone turqueti* samples at a circumpolar level (n = 93) | |
| --- | --- |
| Overall model fit |  |
| Model deviance | 13.779 |
| Percentage explained | 15.418 |
| p-value | <0.001 |
| Relative parameter importance (p-value) |  |
| Geographic distance | 39.847 (<0.001) |
| Sample collection depth | 51.718 (0.057) |
| **(B)** *Pareledone turqueti* samples across the Scotia Sea (n = 52) | |
| Overall model fit |  |
| Model deviance | 2.55 |
| Percentage explained | 37.606 |
| p-value | <0.001 |
| Relative parameter importance (p-value) |  |
| Geographic distance | 38.401 (<0.001) |
| Sample collection depth | 1.010 (0.552) |

Data S1. (separate file: DataS1-S2.xlsx)

Sample information of *Pareledone turqueti* (n = 96) sequenced with target capture sequencing of ddRAD loci.

Data S2. (separate file: DataS1-S2.xlsx)

Sample information of all Southern Ocean octopod samples (n = 440, including 22 technical replicates) sequenced with ddRAD sequencing.

**References (continued from Main Text)**

67. M. C. Fitzpatrick, K. Mokany, G. Manion, M. Lisk, S. Ferrier, D. Nieto-Lugilde, gdm: Generalized Dissimilarity Modeling. R package version 1.4.2.2. (2021).

85. A. Canty, D. Ripley, boot: Bootstrap R (S-Plus) Functions. R package (2022).
